# Quantitative analysis of MHC class II peptide exchange reveals pivotal role of peptide association rate

**DOI:** 10.1101/2024.02.12.579932

**Authors:** Matthias Günther, Jana Sticht, Christian Freund, Thomas Höfer

## Abstract

MHC-II presents antigenic peptides to T helper cells, thus shaping adaptive immune responses. Peptide loading of MHC-II in endosomes is shaped by the susceptibility of the peptide-MHC-II complex to dissociation by the catalyst HLA-DM. For a given MHC-II allotype, experimental data reveal an enormous range of HLA-DM susceptibilities of different peptides – more than five orders of magnitude. To understand the underlying mechanisms, we develop a coarse-grained kinetic model and confront it with experimental data. The model explains the observed variation of HLA-DM susceptibility with the peptide-MHC-II binding energy by an allosteric competition mechanism. Paradoxically, however, certain peptides are resistant to dissociation by HLA-DM regardless of their binding energy. Our model predicts that this resistance is linked with fast peptide association to MHC-II in the absence of HLA-DM. In sum, our data-based theoretical analysis identifies two distinct molecular mechanisms that shape antigen presentation by MHC-II.

## INTRODUCTION

The class II major histocompatibility complex (MHC-II) is expressed on professional antigen-presenting cells and presents peptides derived from endocytosed proteins to CD4 T cells. MHC-II is loaded with these peptides in the late endosomal compartment, from where peptide-MHC-II (pMHC-II) complexes are transported to the cell surface. During peptide loading, an endogenous peptide, class II-associated invariant chain peptide (CLIP), is exchanged against endosomal antigens. This process requires the exchange catalyst human leukocyte antigen DM (HLA-DM). The mixture of antigenic peptides can be very diverse and the MHC-II proteins themselves are highly polymorphic, which poses challenges to accurately predicting which peptides are presented by which MHC-II allotypes [1]. Recent research has provided detailed structural insights into the interactions between peptide-MHC-II and HLA-DM [2–6], and molecular dynamics studies are beginning to characterize conformational ensembles [7, 8]. However, this mechanistic understanding has not yet translated into a quantitative, systems-level model of the HLA-DM-catalyzed peptide switch that relates the biophysical properties of peptides to their presentation by MHC-II [9].

Despite their high degree of polymorphism, MHC-II proteins have a common fold—they are αβ-heterodimers that form a slightly curved β-sheet as the bottom of the peptide binding groove, and two α-helices, one from each chain, lining the groove. The peptide binding groove sits on top of two membrane-proximal immunoglobulin-like domains, one from each chain, and transmembrane helices anchor both chains to the membrane. While the overall fold is well defined, MHC-II proteins are marked by structural plasticity, in particular in the region of the binding groove interacting with the N-terminal part of the peptide where the α-helix of the β-chain can partially cover the β-sheet [10]. Indeed, free MHC-II has been shown to exhibit peptide non-receptive and receptive forms, with transition kinetics between these two forms on the time scale of hours [10, 11]. Peptides are held in the peptide binding groove by two sets of anchor points (Fig. 1A): (i) a set of hydrogen bonds between the peptide backbone and the residues located in both α-helices, and (ii) and set of defined contact points—P1, P4, P6, P7, and P9—distributed along the β-sheet interacting with the peptide side-chains. The peptide binding groove thus directly interacts with the peptide along nine residues, however, since both ends of the groove are open, peptides of (in principle) arbitrary length can be bound, with typical lengths ranging from 10 to 25 residues.

**Figure 1:**
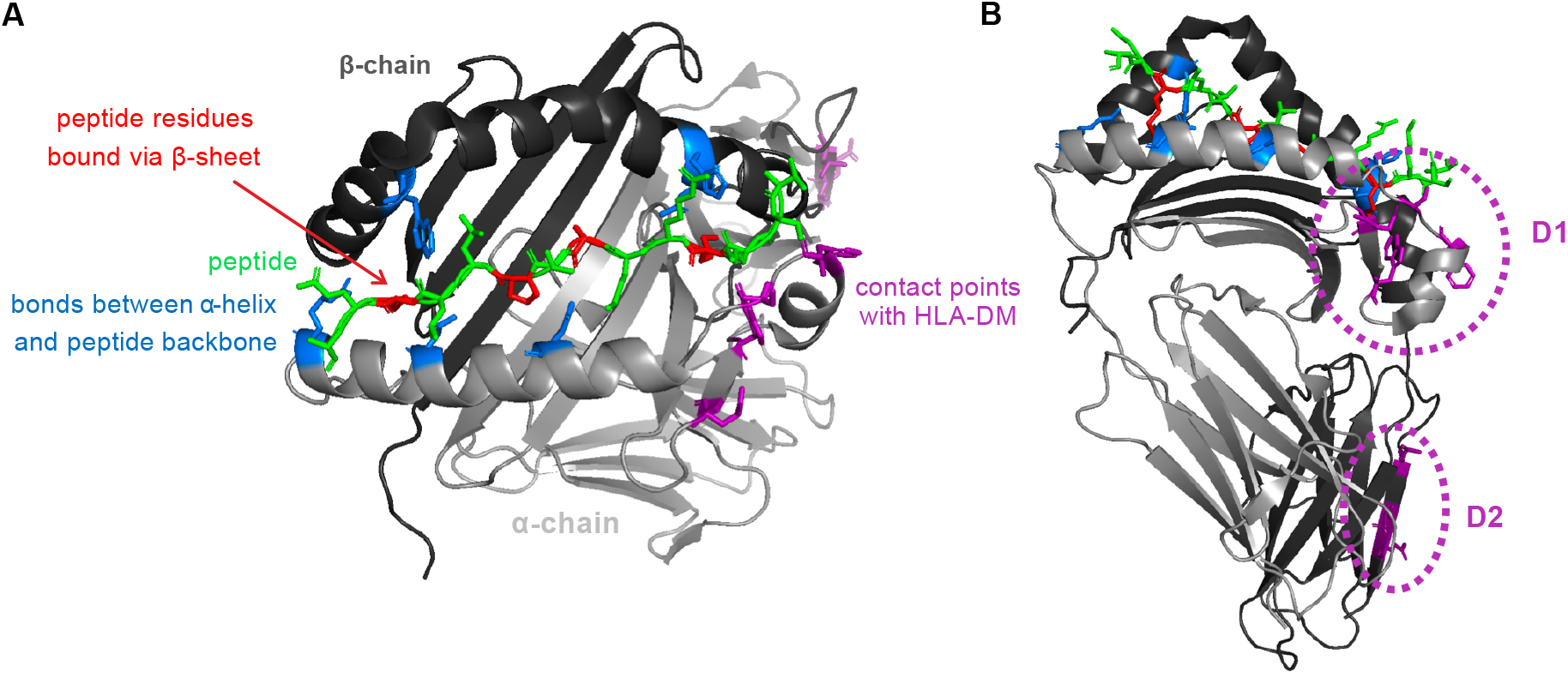
Anchor points on MHC-II proteins for peptide and HLA-DM binding. (A) Peptide-loaded binding groove of MHC-II proteins. The peptide (green) is held by a set of unspecific anchor points (blue) formed between the α-helices of MHC-II and the backbone of the peptide, and a set of specific anchor points formed between the β-sheet and key residues of the peptide (highlighted in red). (B) The D1 and D2 subdomains of the HLA-DM binding domain.

The structure and sequence of HLA-DM is very similar to that of MHC-II proteins, although HLA-DM lacks the ability to bind peptides due to the absence of a peptide binding groove [12]. The HLA-DM binding domain of MHC-II proteins comprises several anchor points that can be divided into two groups [13]: the main anchor points are located on the α-chain near the N-terminal peptide binding groove (D1 subdomain, Fig. 1B), and a second group of weaker anchor points is located in the immunoglobulin-like domain of the β-chain near the transmembrane region (D2 subdomain, Fig. 1B). While it was originally believed that HLA-DM forms long-lived complexes of high affinity with free MHC-II newer findings suggest a rather short-lived and unstable MHC-II-HLA-DM complex. Nevertheless, HLA-DM enhances peptide association already at nM concentrations [11]. Moreover, HLA-DM can bind to pMHC-II complexes, which destabilizes the complex and enhances peptide dissociation [14]. Taken together, the actions of HLA-DM cause particularly stable pMHC-II complexes to be favored for antigen presentation.

HLA-DM shapes the immunopeptidome by catalyzing MHC-II loading differentially for distinct peptides. To characterize this property, Reyes-Varges et al. used a real-time fluorescence anisotropy assay to measure kinetic parameters of HLA-DM enzymatic activity for a range of peptides that exhibit distinct intrinsic stabilities [14]. Intrinsic stability predominantly depended on the peptide side chains that interact with the N-terminal peptide binding groove. Indeed, for the used MHC-II allotype, DRB1^*\**^01:01, the peptide amino acid that contacts P1 has the largest impact on intrinsic stability, while the peptide amino acids interacting with the C-terminal peptide binding groove are less influential (while for other allotypes P4 is the most defining pocket). Interestingly, the authors found that several peptides have a high tendency to resist HLA-DM activity. Both highly stable and less stable pMHC-II complexes were found to exhibit HLA-DM resistance. Together, these findings suggest that both intrinsic stability of peptide binding and HLA-DM resistance play a key role in peptide selection for the immunopeptidome.

To understand the structural basis of peptide exchange, Wieczorek et al. performed molecular dynamics simulations of the exchange pathway finding intermediate states along the exchange pathway [7]. In particular, an excited state provides a candidate for how HLA-DM binding destabilizes the pMHC-II complex. However, the quantitative interplay of multiple conformational mechanisms on relevant timescales for pMHC-II stability of minutes to months is difficult to analyze with molecular dynamics simulations. To address this fundamental question, we here develop a coarse-grained description of the interactions between MHC-II, peptide and HLA-DM that bridges between conformational states and the “macroscopic” dynamics that determine the measurable population frequencies of the various complexes. Within this framework, we gain analytical insight into the parameters that govern peptide loading of MHC-II and derive experimentally testable predictions. Unexpectedly, we find that the association rate of a peptide with MHC-II governs its resistance against HLA-DM-catalyzed release.

## RESULTS

### Modeling the population dynamics of HLA-DM-catalyzed peptide exchange

We begin by developing a basic mathematical framework that links the quantitative mechanisms of HLA-DM-catalyzed peptide exchange with experimental observations. Peptide and HLA-DM can bind with 1:1 stoichiometry to MHC-II proteins, while peptides and HLA-DM do not interact directly with each other. Hence, this stoichiometric interaction model comprises six macroscopic reaction species: free MHC-II, free HLA-DM, free peptide, the binary MHC-II-HLA-DM complex, the binary pMHC-II complex, and the ternary peptide-MHC-II-HLA-DM complex (Fig. 2A). These six reaction species interconvert via four reversible, bimolecular binding reactions: (i) basal peptide binding, (ii) basal HLA-DM binding, (iii) HLA-DM-catalyzed peptide binding, and (iv) peptide-modulated HLA-DM binding. Each of these reactions is characterized by an on-rate and an off-rate. The values of these rate constants depend on the MHC-II allotype and the peptide sequence, which will account for allotype-specific antigen presentation profiles. The ultimate aim of our theory is to explain the peptide-specific on- and off-rates from the molecular properties and dynamics of the microscopic binding domain (Fig. 1).

**Figure 2:**
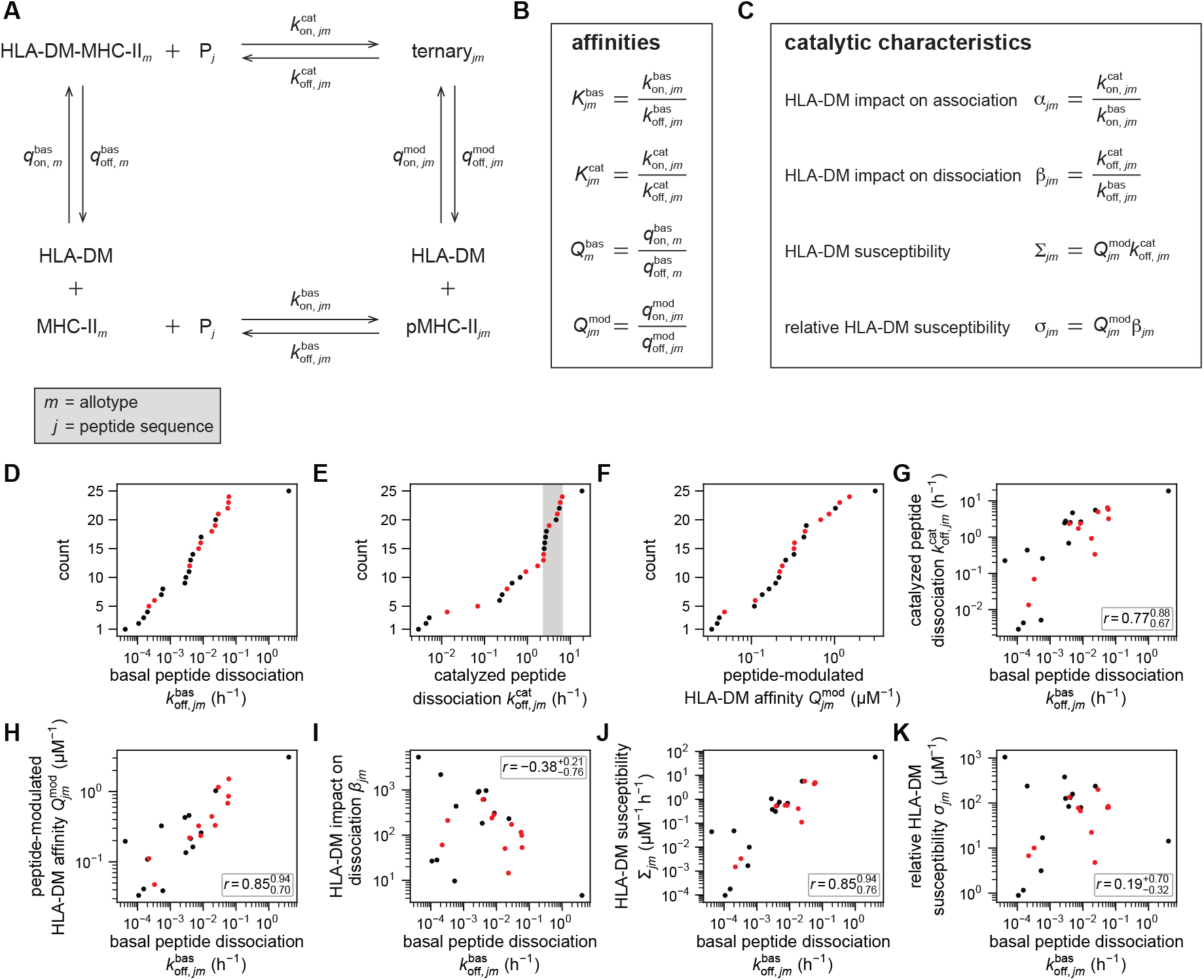
Stoichiometry-based, mathematical model and experimental data. (A) Principal interactions between MHC-II proteins of allotype *m*, peptides of sequence *j* (*P*_*j*_), and HLA-DM, which depend on eight macroscopic rate constants (reaction arrows), which give rise to (B) affinities and (C) catalytic characteristics. (D)-(F) Experimental data of (D) basal peptide dissociation rate 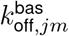, (E) catalyzed peptide dissociation rate 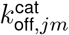, (F) and peptide-modulated HLA-DM affinity 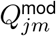 for various peptides and the same allotype, DRB1^*\**^01:01. (G)-(K) Relations between catalytic parameters and basal peptide dissociation rate; r-values are Pearson’s correlation coefficients of the log-values shown with 95% confidence intervals obtained by non-parametric bootstrap. The peptide set contains highly related peptides (black dots) and unrelated peptides (red dots), cf. Fig. S1. Data taken from [14].

Specifically, we introduce an MHC-II allotype index *m* and a peptide sequence index *j*. Thus, each pair of MHC-II allotype and peptide (*jm*-pair) is characterized by eight rate constants (Fig. 2A), where the on-rate and off-rate of basal HLA-DM binding depend only on the MHC-II allotype *m* while the remaining six rate constants depend also on peptide sequence *j* (in general, whether a parameter depends on the MHC-II allotype and/or peptide sequence is indicated in the subscript of the parameter name). Detailed balance imposes a constraint on the affinities of the four reactions (Fig. 2B),

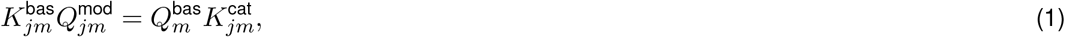

leaving us with seven independent parameters. Two of these (e.g., 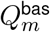 and 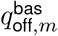) are only allotypespecific while five (e.g., 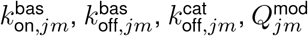 and 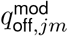) are also peptide-specific.

To characterize the impact of HLA-DM on reversible peptide binding experimentally, often the HLA-DM susceptibility of a pMHC-II complex is evaluated. Susceptibility can be understood systematically by considering the effective peptide association and dissociation rates, 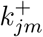, respectively 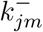,

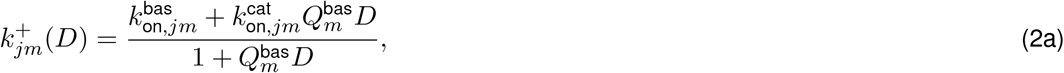

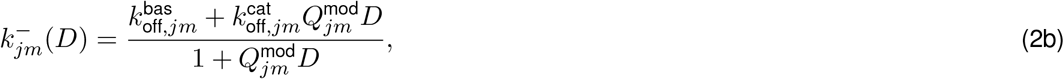

which are both saturating functions of the HLA-DM concentration *D*. Susceptibility,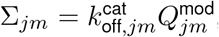, is the linear-response term of effective peptide dissociation, describing the change of 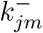 with HLA-DM concentration when HLA-DM concentration is low,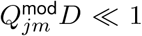. Eqs. (2) suggest to define further catalytic characteristics of HLA-DM action (Fig. 2C). HLA-DM-impact on association 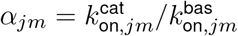, respectively HLA-DM-impact on dissociation 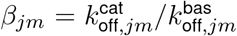, describe the fold-change of the corresponding rates between the absence of HLA-DM, *D* = 0, and saturating concentrations, *D → ∞*. Finally, we consider the relative susceptibility, 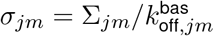. This parameter provides a fundamental link between peptide dissociation and peptide association,

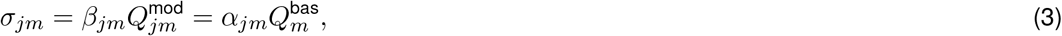

where the first equality follows from the definition of HLA-DM susceptibility, and the second equality follows from detailed balance. This relation will play a central role for model analysis. In summary, the stoichiometric interaction model defines observable parameters—rate constants, affinities, and catalytic characteristics—that determine the population-level dynamics of an MHC-II peptide pair.

### Catalyzed peptide exchange is shaped by peptide-unspecific interactions

With this framework at hand, we turn to experimental observations. In a recent study, three of the peptide-specific parameters — 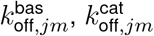 and 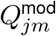 — were determined for a set of 25 peptide sequences, all in the context of the same MHC-II allotype, DRB1^*\**^01:01 [14]. Out of the three parameters, basal peptide dissociation is the most variable, spanning a little more than five orders of magnitude, with the lifetimes ranging from a few minutes up to three years (Fig. 2D). The second most variable parameter is the catalyzed peptide dissociation rate, with variation spanning a little more than three orders of magnitude and the lifetimes ranging from a few minutes up to two weeks (Fig. 2E). The least variable parameter of the three is peptide-modulated HLA-DM affinity spanning not quite two orders of magnitude, with the lowest affinity being 30 µM and the highest 320 nM (Fig. 2F). Taken together, the data cover a broad spectrum of dynamical behavior, and the patterns found within the data can thus be expected to be informative on mechanistic determinants of the interactions between peptide, MHC-II and HLA-DM.

An immediately noticeable pattern is that 12 out of 25 peptides exhibit very similar values in catalyzed peptide dissociation rate of about 5 h^*−*1^ (Fig. 2E, gray area). This observation is remarkable when looking at the sequence similarity of the peptides. 14 out of 25 peptides are from similar sequences, related by single-amino-acid substitutions (Fig. S1A; black dots in Fig. 2D-K), whereas the remaining 11 peptides are unrelated (Fig. S1B; red dots in Fig. 2D-K); the peptides with common catalyzed dissociation rates come from both subsets (Fig. S1, marked by a star). This observation suggests that peptide-unspecific interactions, between the peptide backbone and the α-helices of the binding groove, determine an upper limit for catalyzed peptide dissociation rate. Peptide-unspecific interactions could also play a role for catalyzed peptide association, as measurements of the catalyzed association rates of two distinct peptides (the placeholder peptide CLIP and influenza-derived peptide HA) have yielded very similar values [11]. Thus, peptide-unspecific interactions appear to play a key role in both HLA-DM-catalyzed peptide association and dissociation.

### Catalytic impact of HLA-DM correlates with peptide-specific dissociation rate

Next, we analyzed the dependence of key parameters on the basal peptide dissociation rate (Fig. 2G-K), as we expect the latter to reflect the binding energy of pMHC-II complexes. The data show that catalyzed peptide dissociation rates tend to increase with basal peptide dissociation rate (Fig. 2G). This suggests that the binding energy of the pMHC-II complex is also an important regulator of the stability of the peptide-MHC interactions when in ternary complex with HLA-DM. Furthermore, peptide-modulated HLA-DM affinity exhibits an increasing trend with basal peptide dissociation as well (Fig. 2H). This suggests that high-energy pMHC-II complexes prevent proper HLA-DM binding, while HLA-DM can only tightly bind MHC-II if the pMHC-II complex has low binding energy. Interestingly, the opposite is observed for HLA-DM-impact on dissociation (Fig. 2A): high-energy pMHC-II complexes tend to have a larger HLA-DM-impact on dissociation than low-energy complexes (Fig. 2I). This suggests that the energy that HLA-DM brings into the system is either used to cause a large change in peptide dissociation rate, and HLA-DM has therefore little energy left to stabilize its own binding (high-energy pMHC-II complexes), or it is used to cause a small change in peptide dissociation rate, and HLA-DM can use most of its energy for stable contact with the HLA-DM binding domain (low-energy pMHC-II complexes). These intriguing patterns in the experimental data raise the question which molecular mechanisms determine this biphasic effect of HLA-DM.

Both catalyzed peptide dissociation rate and peptide-modulated HLA-DM affinity show an increasing trend with basal peptide dissociation rate. Consequently, HLA-DM susceptibility shows a strong, increasing trend with basal peptide dissociation rate as well (Fig. 2J). Interestingly, relative susceptibility appears largely uncorrelated from basal peptide dissociation rate; however, it nevertheless is subject to substantial variation spanning about three orders of magnitude (Fig. 2K). This observation suggests that relative HLA-DM susceptibility is not determined by the binding energy of pMHC-II complexes. Taken together, the data provide valuable input for the model to understand how HLA-DM impacts peptide-MHC-II interactions.

### Competition of HLA-DM and peptide results in two regimes for peptide dissociation

In the remainder of the paper, we ask to what extent the known interactions of peptide, MHC-II and HLA-DM explain the quantitative patterns in the experimental data (Fig. 2G-K). To this end, we will equip the macrostates of the stoichiometric interaction model (Fig. 2A) with inner microstates that reflect the actual molecular interactions and corresponding conformational states (Fig. 1, Supplemental Methods). Previous molecular dynamics simulations of the pMHC-II have indicated the existence of two key conformations, a ground state in which the peptide is strongly bound (Fig. 3A, *X*^*β*^), and an excited state (Fig. 3A, *X*^*α*^), from which the peptide can dissociate and which is prone to DM-catalyzed exchange [7]. In the ground state, MHC-II assumes a conformation optimal to accommodate the peptide (light red), in which the peptide is held by both the unspecific contacts formed between the α-helices of MHC-II and the backbone of the peptide, and the specific anchor points in the β-sheet interacting with the side chains of the peptide. In order for the peptide to leave the binding groove, it must pass the α-helices, suggesting that the binding energy of the excited state is dominated by unspecific contacts of the peptide backbone with the α-helices (the possibility that specific interactions between the peptide side chains and the α-helices also play a role is discussed in a more refined model further below). Consequently, the microscopic dissociation rate from the excited state, 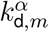, is peptide-unspecific (Fig. 3A, Table 1). Hence, in this approximation, the only peptide-specific parameter of the microscopic model is the free-energy difference between ground state and excited state, which we will refer to as pMHC-II binding energy, denoted 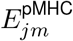. As a result, the measurable basal rate of peptide dissociation 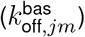 is a function of peptide-specific binding energy and the unspecific dissociation rate from the excited state (Supplementary Methods), demonstrating how the introduction of molecular microstates provides mechanistic underpinning to the stoichiometric parameters. Quantitatively, binding energies between different peptides and MHC-II molecules differ by several tens kJ mol^*−*1^, giving rise to variation of basal peptide dissociation rate over several orders of magnitude (Fig. 3B). Thus, the microstate model naturally accounts for the broad spectrum of observed basal dissociation rates.

**Table 1:** Summary of the microscopic model parameters. The model contains five strictly MHC-II-allotype-specific parameters (subscript *m*) and two parameters that are also peptide-specific (subscript *jm*). The binding domains of MHC-II are defined in Fig. 1.

| parameter | description |
| --- | --- |
| $k_{a,m}^{\alpha}$ | minimal peptide association rate |
| $k_{d,m}^{\alpha}$ | microscopic peptide dissociation rate from the excited state (α-helices) |
| $Q_m^{D2}$ | microscopic affinity of HLA-DM for the D2 subdomain |
| $E_m^{DM}$ | energetic potential of HLA-DM for binding the D1 subdomain |
| $E_m^{\text{bar}}$ | reaction barrier posed by free MHC-II for basal peptide binding and for basal HLA-DM binding of the D1 subdomain |
| $E_{jm}^{\text{pMHC}}$ | binding energy between peptide and the peptide binding groove (α-helices and β-sheet) |
| $x_{jm}$ | variation in basal peptide association |

**Figure 3:**
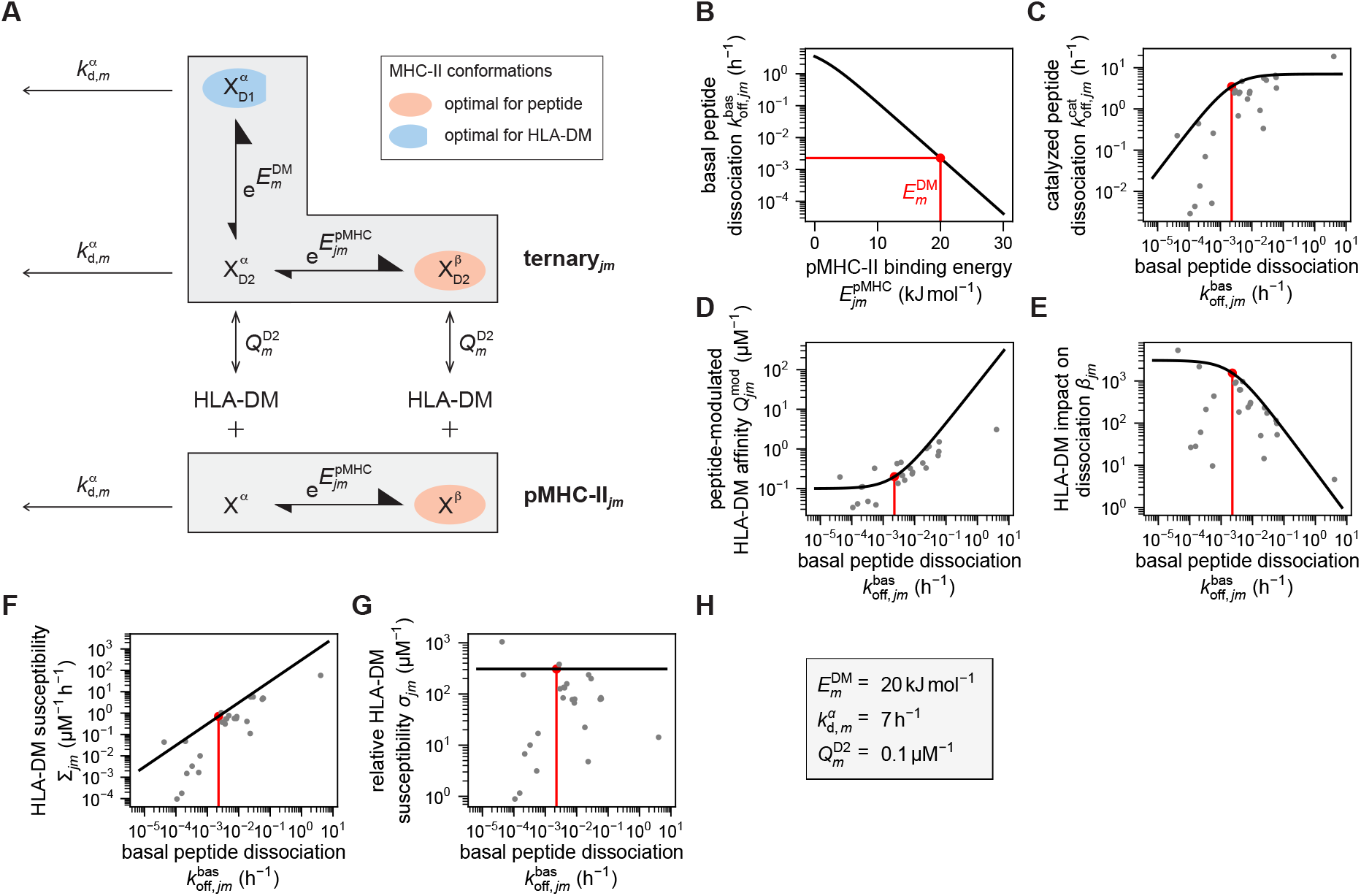
The interplay of pMHC-II binding energy and the binding energy of HLA-DM for MHC-II explains trends in experimental data. (A) Microscopic model for pMHC-II and the ternary complex in which the binding energy 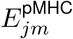 is the only peptide-specific parameter. (B) Basal peptide dissociation rate serves a proxy for pMHC-II binding energy. The energetic potential of HLA-DM binding, 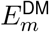, separates weakly and strongly binding pMHC-II complexes (red line). (C)-(G) Comparison between model (black curves) and experimental data (gray dots); the red lines correspond to 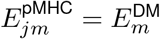. (H) Parameter values used in the model.

Next, we asked how the catalytic action of HLA-DM depends on pMHC-II binding energy. For the ternary complex of pMHC-II and HLA-DM, we now also consider the microstates of HLA-DM interactions with the D1 and D2 subdomains (Fig. 1B, Supplementary Methods). Given the spatial distance to the peptide binding groove, we assume that HLA-DM interacts with the D2 subdomain independent of peptide. By contrast, tight HLA-DM binding, involving also the D1 subdomain, is incompatible with peptide binding in the ground state in which the specific contacts with the β-sheet are made. We denote the energetic potential of HLA-DM binding to the D1 domain by 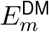. A structural element of MHC-II proteins that may serve as a potential candidate for mediating this competition is MHC-II residue αW43, which needs to be outward-flipped for optimal DM contact [3, 15]. Thus, there are three microstates of the ternary complex, 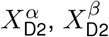 and 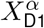 (Fig. 3A). Simultaneously maximizing the binding energies of peptide and HLA-DM is not possible, and only one of the binding partners can assume its preferred configuration with MHC-II at a time (state 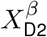 for peptide binding and state 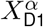 for HLA-DM binding), while the respective other binding partner is forced into a destabilized configuration.

This microstate model (Fig. 3A) allowed us to quantify how catalytic action of HLA-DM depends on pMHC-II binding energy 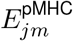 (Table 2). Remarkably, the results account for several principal trends in the experimental data (Fig. 3C-G; note that we used the measurable basal off-rate of the peptide as a proxy for 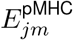, cf. Fig. 3B). Two distinct dynamic regimes emerge: For rather weakly bindig peptides (high 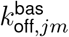), the energetic potential of HLA-DM to the D1 subdomain of MHC-II, 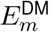, exceeds that of the binding energy of peptide, 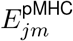, and hence HLA-DM can be stably bound in the ternary complex. Conversely, for strongly binding peptides, we have 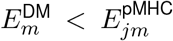 and the entire energetic potential of HLA-DM contributes to destabilize peptide binding without HLA-DM becoming stably bound. The red line in Fig. 3C-G indicates the boundary between the two regimes 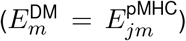. In the high-energy pMHC-II regime, HLA-DM gradually destabilizes the bound peptide (Fig. 3C; linear increase of 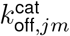 for 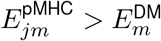) but HLA-DM itself remains weakly bound to MHC-II, mainly interacting with D2 (Fig. 3D; low plateau of the affinity 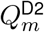). Conversely, in the low-energy pMHC-II regime, peptide binding is dominated by unspecific interactions with the specific interactions with the β-sheet being destabilized by HLA-DM (Fig. 3C; high plateau of 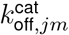 for 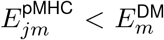), while HLA-DM itself binds more stably by interacting with the D1 subdomain (Fig. 3D; linear increase for 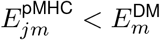). Of note, to reproduce the trends in the experimental data, it was sufficient to adjust three peptide-independent parameters: the energetic potential of HLA-DM to the D1 domain 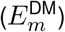, the HLA-DM affinity to the D2 domain 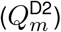 and the peptide dissociation rate from the excited state 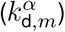 (Fig. 3H). Specifically, the resulting estimate of the HLA-DM binding energy is 20 kJ mol^*−*1^.

**Table 2:** Mathematical formulas of the microscopic model of peptide-MHC-II-HLA-DM interactions, cf. Fig. 4A, in which basal peptide dissociation rate 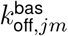 and variation in basal peptide association *x*_*jm*_ act as independent variables (Supplementary Methods). Upper part: kinetic and thermodynamic parameters that appear in the effective peptide binding rates, Eqs (2). Lower part: catalytic characteristics, Fig. 2C. The microscopic model parameters appearing in the formulas are summarized in Tab. 1. In order to keep the notation simple, thermal energy *RT*, with *R* = 8.314 J K^*−*1^ mol^*−*1^ the gas constant and *T* absolute temperature, which appears in the exponential terms, has been absorbed into the energy terms. Optimal HLA-DM catalysis corresponds to *x*_*jm*_ = 1 (Supplementary Methods).

| Parameter | Formula |
| --- | --- |
| basal peptide association | $k_{\text{on},jm}^{\text{bas}} = x_{jm} k_{\text{a},m}^{\alpha}$ |
| catalyzed peptide association | $k_{\text{on},jm}^{\text{cat}} = \frac{(x_{jm} + e^{E_m^{\text{DM}}}) k_{\text{a},m}^{\alpha}}{1 + e^{E_m^{\text{DM}} - E_m^{\text{bar}}}}$ |
| catalyzed peptide dissociation | $k_{\text{off},jm}^{\text{cat}} = \frac{(x_{jm} + e^{E_m^{\text{DM}}}) k_{\text{d},m}^{\alpha} k_{\text{off},jm}^{\text{bas}}}{x_{jm} k_{\text{d},m}^{\alpha} + e^{E_m^{\text{DM}}} k_{\text{off},jm}^{\text{bas}}}$ |
| basal HLA-DM affinity | $Q_m^{\text{bas}} = \left(1 + e^{E_m^{\text{DM}} - E_m^{\text{bar}}}\right) Q_m^{\text{D2}}$ |
| peptide-modulated HLA-DM affinity | $Q_{jm}^{\text{mod}} = \left(1 + \frac{e^{E_m^{\text{DM}}} k_{\text{off},jm}^{\text{bas}}}{x_{jm} k_{\text{d},m}^{\alpha}}\right) Q_m^{\text{D2}}$ |
| HLA-DM impact on association | $\alpha_{jm} = \frac{1}{x_{jm}} \frac{x_{jm} + e^{E_m^{\text{DM}}}}{1 + e^{E_m^{\text{DM}} - E_m^{\text{bar}}}}$ |
| HLA-DM impact on dissociation | $\beta_{jm} = \frac{(x_{jm} + e^{E_m^{\text{DM}}}) k_{\text{d},m}^{\alpha}}{x_{jm} k_{\text{d},m}^{\alpha} + e^{E_m^{\text{DM}}} k_{\text{off},jm}^{\text{bas}}}$ |
| HLA-DM susceptibility | $\Sigma_{jm} = \left(1 + \frac{e^{E_m^{\text{DM}}}}{x_{jm}}\right) Q_m^{\text{D2}} k_{\text{off},jm}^{\text{bas}}$ |
| relative HLA-DM susceptibility | $\sigma_{jm} = \left(1 + \frac{e^{E_m^{\text{DM}}}}{x_{jm}}\right) Q_m^{\text{D2}}$ |

The two regimes for the catalytic action of HLA-DM are also clearly visible when plotting the HLA-DM-impact on dissociation (i.e., the catalyzed dissociation rate relative to the basal one) (Fig. 3E). The HLA-DM susceptibility itself increases continuously as the pMHC-II binding energy drops (Fig. 3F). When normalizing HLA-DM susceptibility with the basal peptide off-rate, yielding the relative susceptibility (Fig. 2C), we find that the trend in HLA-DM susceptibility generated by the model is entirely due to the peptide off-rate (Fig. 3G). Importantly, however, the experimental data for the relative susceptibility are not constant but show considerable, albeit uncorrelated, scatter. This observation implies that, beyond the pMHC-II binding energy, there must be further parameters affecting the HLA-DM susceptibility.

In conclusion, the competition of peptide and HLA-DM for strong binding to MHC-II explains the characteristic trends in the experimental data on peptide dissociation and HLA-DM binding. However, there is still considerable variability in the data whose explanation requires additional molecular mechanisms.

### DM-resistant peptides have large basal association rates

To account for the as yet unexplained variability in the experimental data (in particular, Fig. 3G), additional model parameters must become peptide-specific (Supplementary Methods). As peptide binding itself depends, of course, already on the nature of the peptide (via 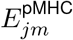), we reasoned that the energetic potential of HLA-DM to the D1 subdomain could additionally show peptide specificity. To formally do this, we introduced a peptide-specific factor 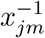 that scales 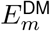 (Fig. 4A). To satisfy detailed balance, we also scale the basal association rate of the peptide with *x*_*jm*_, with intriguing mechanistic consequences that will be discussed below. For this modified microscopic model, we computed the kinetic parameters and catalytic characteristics (Table 2). Remarkably, the simple modification of the energetic potential of HLA-DM generates practically all the additional variability seen in the experimental data (Fig. 4B-F).

**Figure 4:**
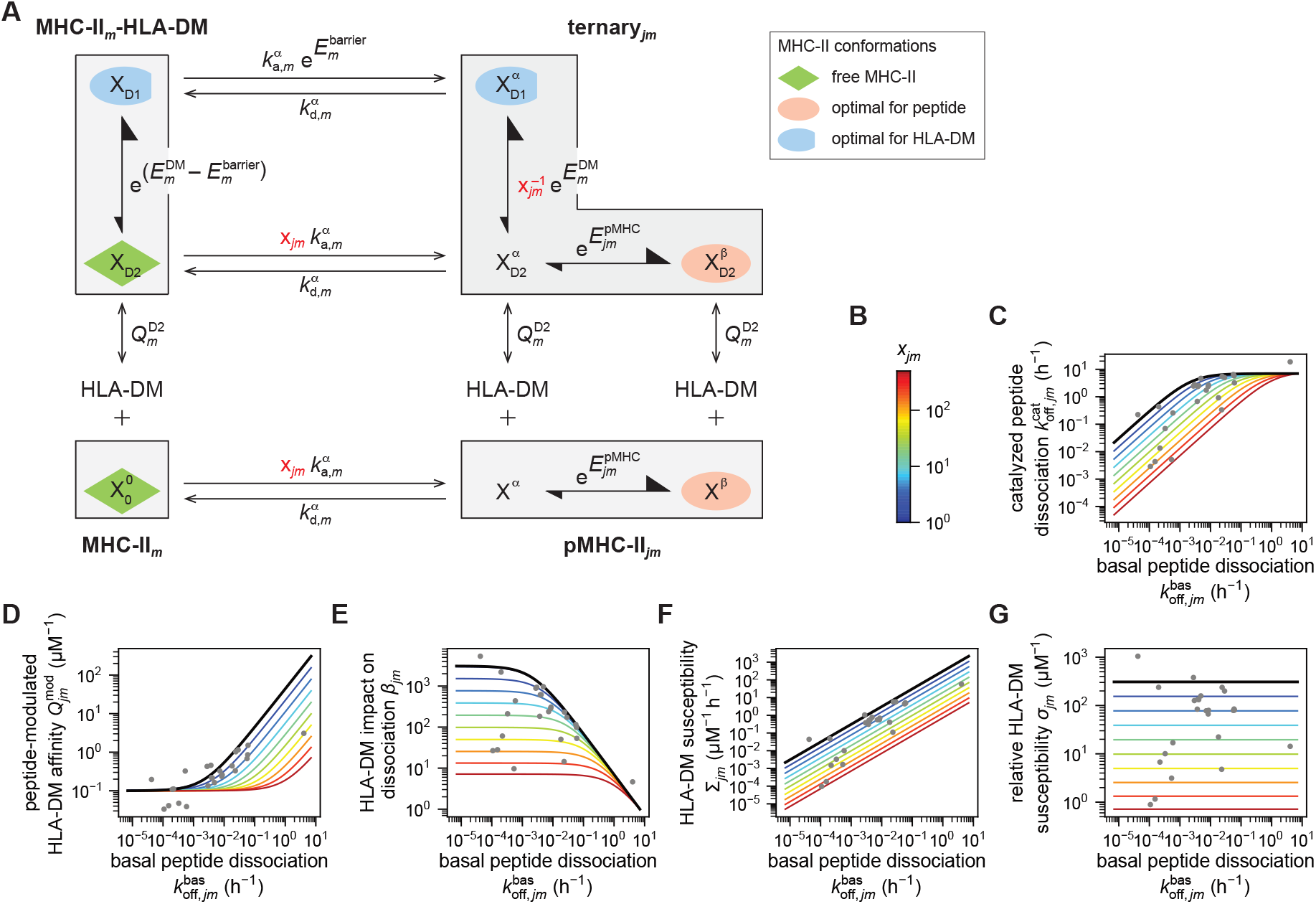
Variation in basal peptide association rate influences the binding energy of HLA-DM for MHC-II. (A) Extension of the microscopic model (Fig. 3A) to include free MHC-II and the HLA-DM-MHC-II complex, in which basal peptide association rate is the second peptide-specific variable, described by the parameter *x*_*jm*_ (highlighted in red). (B) Color bar for the range of values of *x*_*jm*_. (C)-(G) Comparison of the model to the experimental data. The parameter values are the same as in Fig. 3H.

How can this modification of the energetic potential HLA-DM be explained mechanistically? For detailed balance, the peptide-specific factor *x*_*jm*_, could also scale peptide dissociation from the excited state, or peptide association. Closer analysis predicts that solely the peptide association rate is responsible for the HLA-DM resistance seen in the relative susceptibility. To see this, we focus on the large variability of the relative susceptibility in the data (Fig. 4G). We recall that this parameter links peptide dissociation and peptide association, via Eq. (3). Now consider two peptides, *j*_1_ and *j*_2_, binding to the same MHC-II allotype, whose relative susceptibilities have ratio *ρ*. According to Eq. (3), we have

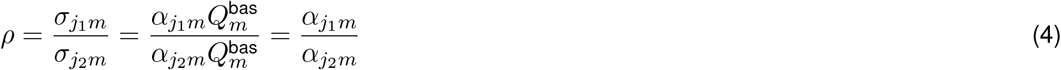

As the basal HLA-DM affinity for peptide-unbound MHC-II 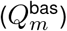 is naturally peptide-unspecific, the ratio of the relative susceptibilities of these peptides is determined by the impact that HLA-DM has on association. This impact is given by the ratio of peptide association rate with HLA-DM bound and without HLA-DM, 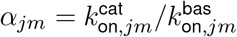 (Fig. 2C). Hence, the variability in relative susceptibility is due to peptide specificity of peptide association.

In principle, this specificity could be due to basal or catalyzed peptide association. As pointed out above, catalyzed peptide association seems to be dominated by peptide-unspecific interactions [11], and hence we make basal association peptide-specific (scaling 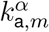 with *x*_*jm*_). The data also suggest that HLA-DM accelerates peptide association, possibly by stabilizing a peptide-receptive conformation of MHC-II [11]. To account for this observation in the model, we introduced a free-energy difference 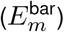 between peptide-bound (i.e., peptide-receptive) and unbound MHC-II (Fig. 4A; without of loss of generality introduced in the catalyzed peptide binding step). The same energy difference must be overcome to bind to the D1 subdomain, which causes the acceleration of peptide association by HLA-DM (Fig. S2). After some algebra, we find that these observation have three implications: (i) the basal HLA-DM affinity is low, (ii) the basal peptide association rate is variable in a peptide-specific manner, and (iii) catalyzed peptide association rate is (approximately) peptide-unspecific. In sum, the peptide specificity of the relative HLA-DM susceptibility (Fig. 4F) is a consequence of the peptide-specific basal association rate. Peptides with low relative HLA-DM susceptibility have high basal association rate, whereas the binding of peptides with high relative HLA-DM susceptibility is poor without HLA-DM and greatly accelerated by HLA-DM.

### Basal peptide association controls the impact of HLA-DM on peptide exchange

Finally, we used the modified microscopic model to predict how HLA-DM activity shapes peptide exchange. To this end, we computed the effective peptide exchange rates (Eqs. (2)) as a function of HLA-DM concentration, *D* (Fig. 5). Generally, HLA-DM accelerates both effective peptide binding and dissociation (Fig. 5A-D). Peptide specificity of the effective association rate is predicted to manifest only at comparatively low HLA-DM concentration, whereas high HLA-DM catalyzes uniformly fast peptide association (Fig. 5E). Conversely, the basal association rate of the peptide is predicted to affect the effective dissociation rate across the entire range of HLA-DM concentration. This effect is qualitatively seen for weakly bound (Fig. 5F), intermediate (Fig. 5G) and strongly bound (Fig. 5H) peptides, and quantitatively most pronounced for strongly bound peptides. Thus, the basal peptide association rate controls the impact of HLA-DM on peptide exchange.

**Figure 5:**
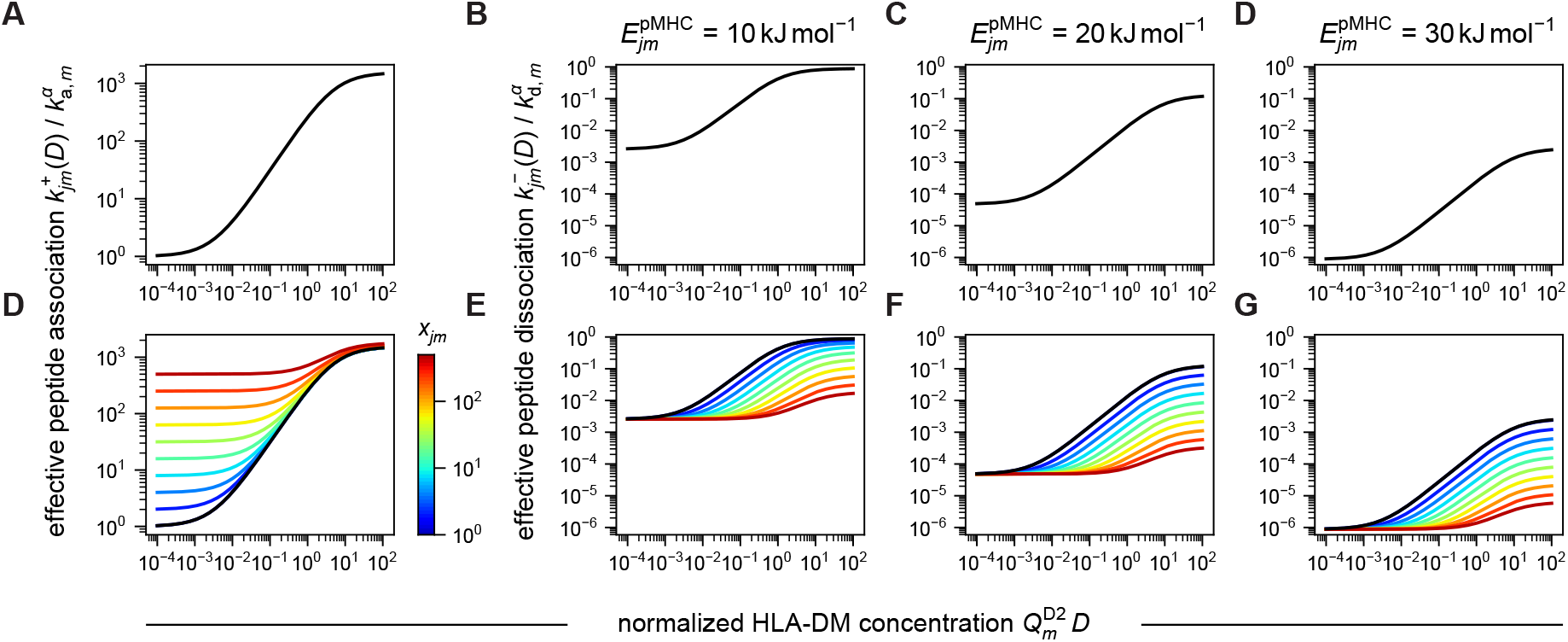
Effective peptide binding rates are strongly influenced by variation in peptide association at all HLA-DM concentrations. (A)-(D) Effective peptide binding in dependence on HLA-DM concentration without variation in basal peptide association rate. (A) Effective peptide association rate and (B)-(D) effective peptide dissociation rate for (B) a weakly bound peptide,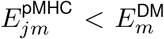, (C) a peptide bound with intermediate binding energy, 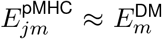, and (D) a strongly bound peptide 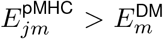 (E) - (H) The same as (A) - (D) but with variation in basal peptide association, parameterized by *x*_*jm*_ (colorbar). The reaction barrier and the energetic potential of HLA-DM are 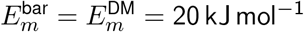.

## DISCUSSION

Here, we derive principle quantitative features of HLA-DM-catalyzed peptide exchange based on known structures and conformational states of the ternary system of MHC-II, peptide and HLA-DM. We find that a competitive effect between HLA-DM and peptide, which prevents both binding partners from simultaneously making optimal contact with MHC-II, results in two distinct regimes of catalyzed peptide exchange. With strongly binding peptides, HLA-DM expends its energetic potential for destabilizing the peptide, leaving comparatively little energy for binding to MHC-II. By contrast, HLA-DM can fully destabilize weakly binding peptides and expend the remaining energetic potential in binding more strongly to MHC-II. Interestingly, with this mechanism, the experimental data predict an HLA-DM energetic potential of *∼* 20 kJ mol^*−*1^. We show that this mechanism implies that the basal dissociation rate of the peptide governs catalyzed peptide exchange.

Importantly, however, the competition mechanism leaves considerable variation in the experimental data unexplained. We find that principal thermodynamic constraints predict the existence of a second, cooperative effect: HLA-DM accelerates the association of the peptide. Hence, the basal peptide association is an independent parameter shaping catalyzed peptide exchange, with rapidly associating peptides being HLA-DM-resistant. These insights can be tested experimentally by measuring peptide association rates. We predict a strong negative correlation between basal peptide association rates and relative HLA-DM susceptibility.

A candidate mechanism realising the cooperative effect could be an allosteric interaction between HLA-DM binding, specifically to the D1 subdomain, and peptide association. Thus, part of the energetic potential of HLA-DM would be expended to stabilize the peptide-receptive conformation of MHC-II. This mechanism implies that there is no stable complex formation between MHC-II and HLA-DM, and predicts that the dissociation constant of this interaction is in the micromolar range.

While the data used to analyze the model were obtained for a single MHC-II allotype (HLA-DRB1^*\**^01:01), we expect the model to apply to other MHC-II allotypes. The question therefore becomes which parameter values may differ/are the same between different MHC-II allotypes. Although MHC-II proteins are highly polymorphic, the HLA-DM binding domain is strongly conserved across MHC-II variants (with the notable exception of HLA-DQ isotypes). This means that the parameters 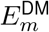 and 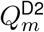, which are associated with the D1, respectively D2 subdomain, could have similar values for different MHC-II allotypes. MHC-II variants predominantly differ in the residues of the β-sheet and, to a somewhat lesser extent, the α-helices of the peptide binding groove. Furthermore, variations tend to be more numerous in the β-chain than in the α-chain (with the most extreme example being the HLA-DR isotype, for which the α-chain is practically monomorphic). The parameters 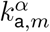 and 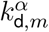, as well as *x*_*jm*_ could be influenced by sequence variations of the α-helices (in particular, basal association could be shaped by interactions of the peptide residues with the α-helices). By contrast, 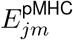 should be sensitive to variations in both the α-helices and the β-sheet. Understanding how MHC-II polymorphism shapes the parameter values remains a key challenge in the systematic quantification of peptide-MHC-II-HLA-DM interactions.

In its current form, the model is appropriate for analyzing dissociation experiments *in vitro*. To test our specific predictions on the role of the peptide association rate, *in vitro* association experiments will be useful. To interpret such experiments, it might be necessary to explicitly incorporate further mechanisms into the model. In particular, free MHC-II can form aggregates in the absence of peptide [11], and accounting for the conservation of monomeric and aggregated MHC-II molecules will be relevant for the quantitative interpretation of association measurements. Both these processes are peptide-unspecific and are relevant for proper quantification of association measurements.

To understand how the processes involved in MHC-II loading *in vivo* qualitatively and quantitatively shape immunopeptidomes, we expect that the current model will become a building block. The physiological situation involves two key factors not studied with the present model: the competition of many different peptides for exchange with the placeholder CLIP, and the non-equilibrium transport of pMHC-II to the cell surface. In principle, the model could predict pMHC-II loading in the presence of a peptide mixture. However, additional mechanisms, such the proposed compare-exchange mechanism [16] may come into play here and the model will need to be extended accordingly. A more principled extension of the model will be the incorporation of the continuous transport of pMHC-II molecules from the endosome, where HLA-DM and low pH shape MHC-II loading, to the cell surface. Here, we expect that both association time and stability of a given pMHC-II complex in the endosome will be selected against the residence time of MHC-II. Based on our prediction that fast-associating peptides (low association time) are also HLA-DM-resistant (high stability of pMHC-II), peptide association rate could be a key parameter shaping immunopeptidomes.

## Supplementary Methods

In this paper, we analyze a microscopic model of peptide-MHC-II-HLA-DM interactions that is based on a detailed description of molecular states of MHC-II proteins interacting with peptide or/and HLA-DM (Figs. 1, 2A and 4A). Specifically, we relate the macroscopic parameters of a simple stoichiometric description of the system to microscopic parameters that account for these detailed state transitions (Tables 1 and 2). In the following, we derive these equations, summarized in Table 2. The kinetic and thermodynamic considerations entering this analysis manifest on two levels. On the one hand, we have the population level, where macroscopic numbers of reactants (MHC-II, peptide, HLA-DM) interact with each other via the formation and decay of complexes. Here the internal states of the reactants are not tracked, defining the purely stoichiometry-based model (Fig. 2A). On the other hand, we have the molecular level. Here, the reactants are complex molecular structures with internal state ensembles and transitions. Hence each molecule or complex is a thermodynamic system itself. Thus, we describe the interactions between peptide, MHC-II and HLA-DM interactions as a thermodynamic system, at the population level, comprising a macroscopic number of thermodynamic systems at the molecular level. To do this efficiently, we assume that the relaxation times on the molecular level are much faster than the relaxation times on the population level. Put differently, we assume the molecular level to be in thermodynamic equilibrium even when the concentrations of the stoichiometry-based reaction species, Fig. 2A, are not (yet) equilibrated. Indeed, the peptide dissociation experiments we analyze occur on population-level time scales of tens of minutes, during which time internal state transition will typically equilibrate.

The equations for these macroscopic rate and equilibrium constants of the stoichiometry-based model describe observable quantities. In order to compare our microscopic model to the experimental data, we consider an important trade-off: the level of detail at which we describe the molecular level versus the information about the molecular level in the population-level data (Fig. 2D-K). Indeed, it will turn out that the complete microscopic model cannot be fully constraint by the available experimental data. Specifically, there are parameters in the microscopic model that only appear in a certain combination in the final equations but never isolated, or drop out in the derivation of these equations. In other words, there are several non-identifiable parameters in the model. Hence, in the following, we will distinguish two levels of model. What we will call the structure-based model describes the interaction system in great detail by incorporating the molecular structures shown in Fig. 1. The second model, referred to as observational model, will reflect only those aspects of the structure-based model about which the data are informative. The microscopic model shown in Fig. 4A is of the second kind, and hence describes the system as seen in the experiment. In the following, we derive the structure-based model and then show how it gives rise to the observational model of Fig. 4A.

This analysis is organized as follows. First, we derive the structure-based model for the special case that we term of optimal HLA-DM catalysis (Fig. S3). In this version, HLA-DM can use its entire binding potential to destabilize peptide binding to MHC-II, and, at the same time, maximally accelerate peptide association. Furthermore, the optimal-HLA-DM-catalysis model is characterized by a single peptide-specific parameter—the pMHC-II binding energy 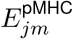—which encodes for the basal peptide dissociation rate. We then discuss the non-identifiabilities and how these give rise to the observational model of optimal HLA-DM catalysis, which corresponds to the model shown in Fig. 3A, and, when setting *x*_*jm*_ = 1, to the model shown in Fig. 4A. As the final part of this analysis, we consider two deviations from optimal HLA-DM catalysis, imperfect peptide-HLA-DM cooperativity (Fig. S4) and imperfect peptide-HLA-DM competition (Fig S5), and discuss the additional peptide-specific parameters that this brings about. One of these deviations, imperfect peptide-HLA-DM cooperativity, gives rise to the observational model of Fig. 4A (where in general *x*_*jm*_ *≠* 1) and represents the main model of this work.

As a last point, we want to emphasize a notational simplification we use throughout the text. Since we conduct a thermodynamic analysis, we frequently encounter Boltzmann factors, i.e., exponential functions in which the arguments are (free) energies *E* (or *G*) divided by the thermal energy *RT*, where *R* = 8.314 J K^*−*1^ mol^*−*1^ is the gas constant and *T* is absolute temperature. In order to avoid too much cluttering of the formulas, we will not explicitly write *RT* in such expressions, e.g., instead of *e*^*G/RT*^ we will simply write *e*^*G*^.

### Optimal HLA-DM catalysis

#### Free MHC-II

We begin with the description of free MHC-II. A key dynamical aspect of MHC-II molecules is that they exhibit structural plasticity. That is, instead of being a rigid body in which each atom has a defined equilibrium position (relative to some co-moving reference frame), certain structural elements exhibit some extent of translational and rotational degrees of freedom, giving rise to a poorly defined fold, which is particularly pronounced in the part of the peptide binding groove interacting with the N-terminal end of the peptide and in the D1 subdomain. To formally handle this, we introduce a parameter *µ* that describes the MHC-II conformation. However, we are not able to give a precise, quantitative characterization of what “conformation” entails, i.e., what the translational and rotational degrees of freedom are that define a conformation and what values they might assume. In the course of this analysis, we rather state the qualitative aspects of the conformations that are relevant for our model. In any case, each MHC-II conformation comes with a free-energy value, 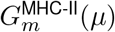, which formally defines the free energy of the MHC-II molecule as a function over the conformation space (free-energy landscape). We assume that there is an MHC-II conformation that minimizes the free energy, and we denote this conformation with 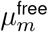, and the corresponding free energy with

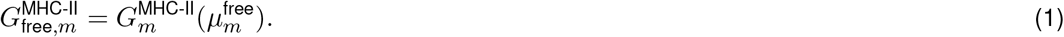

MHC-II conformations obeying 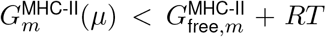 can be visited by free MHC-II due to its interactions with the thermal environment. This set of conformations encompasses the translational and rotational degrees of freedom that lead to free MHC-II molecules with a poorly defined fold.

In order to keep things simple, we nevertheless refer to 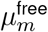 as the conformation of free MHC-II in the following, and we describe the macrostate of free MHC-II as a one-state system, where we denote the corresponding microstate with 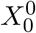 (Fig. S3, MHC-II, green MHC-II conformation). (The distinction between macrostate, microstate, and conformation is somewhat redundant for free MHC-II but it will become relevant in the context of the complexes.) On the population level, the relation between the concentration of free MHC-II, [MHC-II], and the concentration of microstate 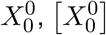 is simply,

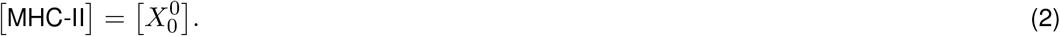

For the MHC-II conformation 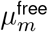 we assume that it has two key characteristics regarding its ability to interact with peptide and HLA-DM. Firstly, the peptide binding groove is “collapsed” and a peptide trying to bind MHC-II must shape the groove upon entering to allow its accommodation. This structural rearrangement increases the free energy of the MHC-II molecule and must be compensated by the binding energy between peptide and the peptide binding groove. Moreover, the association rate of peptide-MHC-II binding can be expected to be small (compared to the hypothetical scenario in which the peptide binding groove of free MHC-II is not collapsed but already optimally shaped for accommodating the peptide). Secondly, the D1 subdomain is “distorted” having its anchor points unfavorably arranged relative to the corresponding anchor points located on the HLA-DM molecule. In order to establish optimal contact with the D1 subdomain, HLA-DM must enforce a structural rearrangement of the MHC-II molecule. This also increases the free energy of the MHC-II molecule, which must be compensated by the binding energy between HLA-DM and the D1 subdomain. A key assumption of our model is that these two structural arrangements are allosterically coupled, which, as we will see in the section on the ternary complex, is the cause behind HLA-DM accelerating peptide association.

### MHC-II-HLA-DM complex

The interactions between MHC-II and HLA-DM are mediated via the two subdomains D1 and D2 (Fig. 1B). This suggests that there are three microstates in the MHC-II-HLA-DM complex (Fig. S3, MHC-II-HLA-DM): HLA-DM is bound (i) only via the D2 subdomain, microstate *X*_D2_, (ii) only via the D1 subdomain, microstate *X*_D1_, or (iii) via both subdomains, microstate *X*_D1/2_. Moreover, HLA-DM may bind to and dissociate from the MHC-II molecule either via the D1 subdomain or the D2 subdomain. We first discuss the binding energies and MHC-II conformations of each microstate. It is assumed that when HLA-DM is bound only via the D2 subdomain, the conformation of the MHC-II molecule remains the same as in its free form (Fig. S3, green MHC-II conformation of microtate *X*_D2_). Denoting the binding energy between the D2 subdomain and the corresponding binding domain on the HLA-DM molecule with 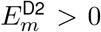, the free energy of the microstate *X*_D2_ is given by

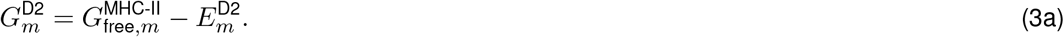

From this microstate, HLA-DM may make further contact to the D1 subdomain, becoming bound via D1 and D2, leading to microstate *X*_D1/2_. As pointed out above, the MHC-II conformation 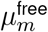 has the D1 subdomain unfavorably arranged for HLA-DM contact. We denote the MHC-II conformation with optimal arrangement of the anchor points of the D1 subdomain with 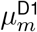 (Fig. S3, light blue MHC-II conformation) and its free energy with 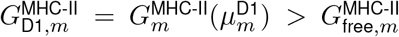. We further denote the binding energy between HLA-DM and MHC-II in conformation 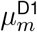 with 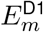, The free energy of microstate *X*_D1/2_ is then given by

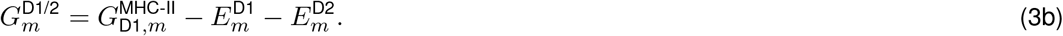

From this microstate, HLA-DM may lose contact with the D2 subdomain, remaining only bound via the D1 subdomain, leading to microstate *X*_D1_. It is assumed that the MHC-II molecule remains in conformation 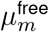, and the free energy of microstate *X*_D1_ is

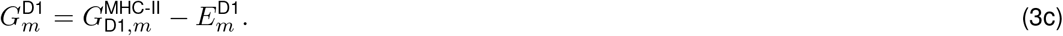

The equilibrium constants of the MHC-II-HLA-DM complex show in Fig. S3 depend on the free-energy differences between microstates. Specifically, the equilibrium constant of transition *X*_D1_ *↔ X*_D1/2_ is determined by

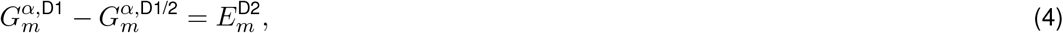

and the equilibrium constant of transition *X*_D2_ *↔ X*_D1/2_ is determined by

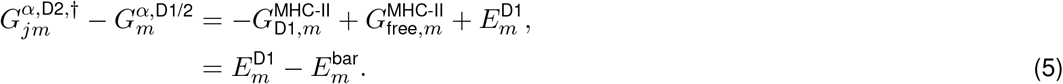

In a last step, we introduce the reaction barrier 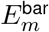, which is defined as the free-energy difference between the MHC-II conformations 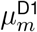 and 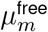,

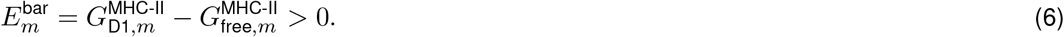

We will later see that this parameter controls the extent by which HLA-DM can accelerate peptide association.

Having determined the free energies of the microstates of the MHC-II-HLA-DM complex, we now consider the population frequencies of these microstates within the complex. On the population level, the concentration of MHC-II-HLA-DM complexes relate to the concentrations of the microstates by

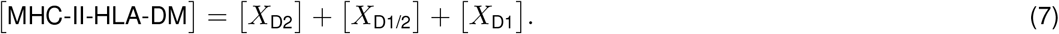

Due to the assumption of rapid molecular relaxation times, the MHC-II-HLA-DM complex is assumed to be in thermodynamic equilibrium. Thus, it holds

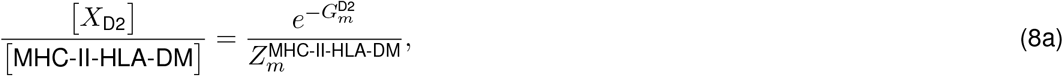

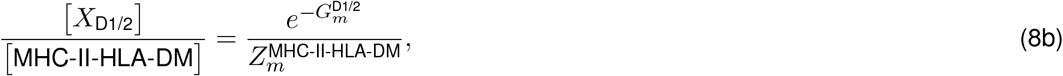

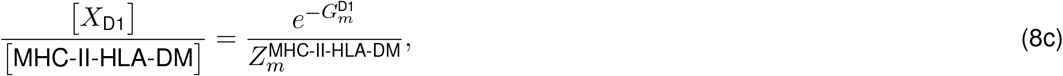

where we have introduced the partition function of the MHC-II-HLA-DM complex,

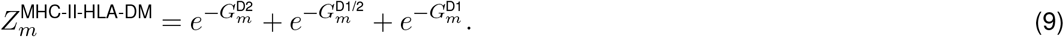

Relations (8) hold even when free MHC-II, free DM and the MHC-II-HLA-DM complex are not in equilibrium on the population level.

Next, we determine the basal HLA-DM affinity as a function of the microscopic parameters. The basal HLA-DM affinity is defined by

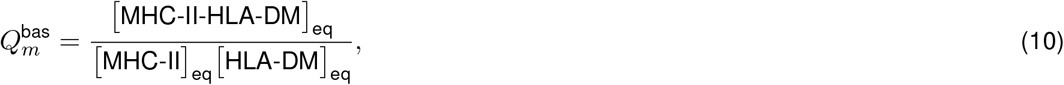

where the subscript eq refers to concentration values in thermodynamic equilibrium on the population level. Microscopically, the MHC-II-HLA-DM complex can form and decay either via the D1 or the D2 subdomain. The corresponding microscopic affinities are defined by

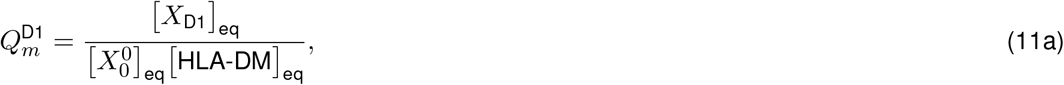

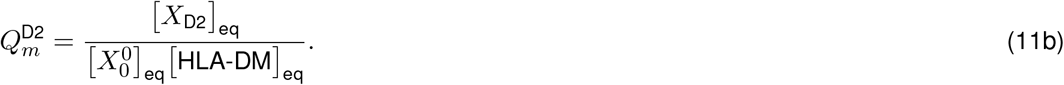

The microscopic affinities obey a detailed-balance relation,

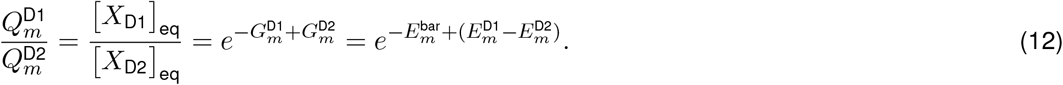

Using relations (2), (6), (8) and definition (11b), we can derive the equation for the basal HLA-DM affinity,

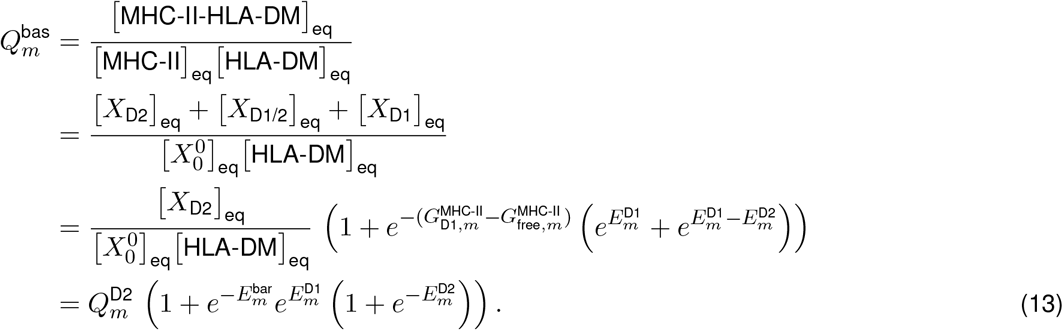

#### pMHC-II complex

The pMHC-II complex is described as a two-state system, comprising the ground state, *X*^*β*^, in which the peptide is well-anchored, and an excited state, *X*^*α*^, from which dissociation occurs and via which the complex forms (Fig. S3, pMHC-II). As outlined above, the conformation of free MHC-II has a “collapsed” binding groove and in order for the peptide to bind, MHC-II has to undergo a structural rearrangement.The MHC-II conformation assumed in the ground state of the pMHC-II complex is denoted 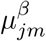 (Fig. S3, light red MHC-II conformation). In this conformation, the peptide binding groove is shaped to allow for optimal interactions between the peptide side chains and the β-sheet of the groove. Furthermore, the key anchor points of the D1 subdomain are not accessible for HLA-DM binding—a property that is one of the two defining features of optimal HLA-DM catalysis and which will become important when discussing the ternary complex. The free energy of MHC-II conformation 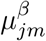 is 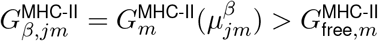. Denoting the overall binding energy between the peptide and the peptide binding groove—comprising contributions from both the interactions between the peptide side chains and the β-sheet of the groove as well as interactions between the peptide backbone and the α-helices of the groove—with 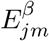, the free energy of the ground state is given by

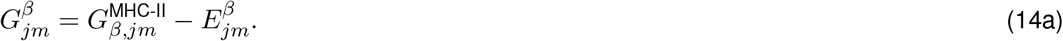

While the binding-energy contribution in the ground state is dominated by peptide-specific interactions, it is assumed that in the excited state peptide-unspecific interactions dominate. Hence, the binding energy of the excited state is taken to be 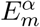, which does not contain the peptide index *j*. (The possibility that this binding energy has relevant contributions from peptide-specific interactions is discussed in Section “Variations beyond optimal HLA-DM catalysis”.) The excited state of pMHC-II lies between the state of free MHC-II, 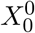, and the ground state *X*^*β*^. The transition from free MHC-II to the ground state of pMHC-II (complex formation) induces a structural rearrangement of the MHC-II molecule from conformation 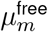 to conformation 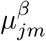 (and the other way around when transitioning from the ground state to free MHC-II, i.e., complex decay). We assume that during complex formation, respectively complex decay, the MHC-II molecule transiently passes through the MHC-II conformation 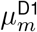, in which the D1 subdomain is optimally arranged for HLA-DM binding (Fig. S3, light blue MHC-II conformation of the excited state). (This is the second defining properties of optimal HLA-DM catalysis, and deviations from this condition will lead to HLA-DM-resistant peptides, which is the topic of Section “Imperfect peptide-HLA-DM cooperativity”.) The free energy of the excited state is given by

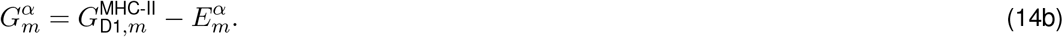

The equilibrium constant for the transition *X*^*α*^ *↔ X*^*β*^ is determined by the free-energy difference between the excited state and the ground state,

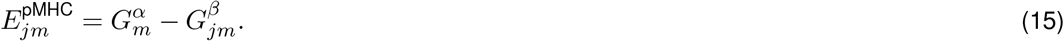

which we refer to as the pMHC-II binding energy.

We now consider the population frequencies of the microstates of the pMHC-II complex. On the population level, the concentration of pMHC-II relates to the concentrations of the two pMHC-II microstates by

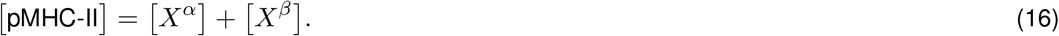

We again employ the assumption of fast relaxation times on the molecular level, and the population frequencies of the microstates become

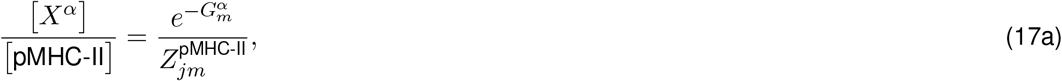

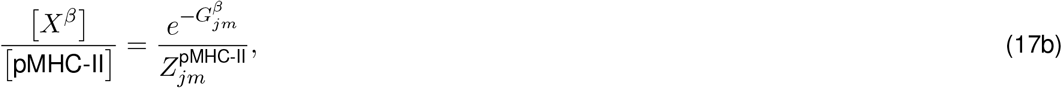

where we have introduced the partition function of the pMHC-II complex,

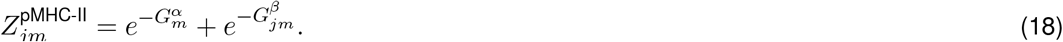

Relations (17) even hold when free MHC-II, free peptide and the pMHC-II complex are not (yet) in thermodynamic equilibrium on the population level.

The kinetics for the transition from free MHC-II, i.e., state 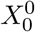, to the excited state are described by mass-action kinetics,

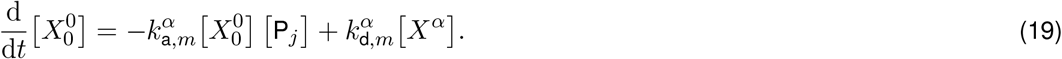

Due to the assumption that the excited state is dominated by peptide-unspecific interactions, the microscopic rate constants 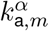 and 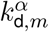 do not depend on the peptide sequence *j*. Using Eqs. (2), (15), (16), (17a) and (18), we arrive at

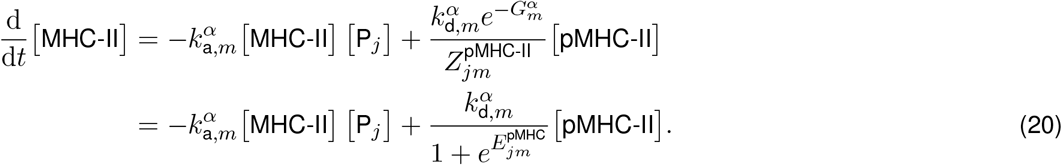

This relation should be compared to the corresponding equation of the stoichiometry-based interaction model,

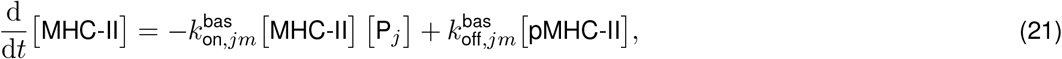

from which we immediately see that

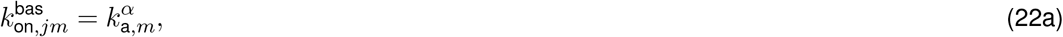

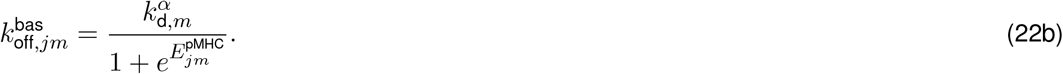

The basal peptide affinity is given by

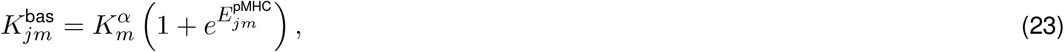

where we introduced the microscopic peptide affinity for the excited state,

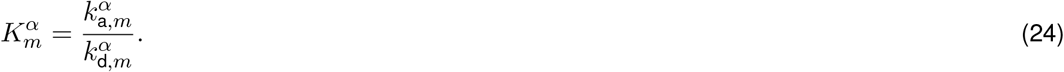

### Ternary complex

Given that the MHC-II-HLA-DM complex contains three microstates and the pMHC-II complex contains two microstates, it seems reasonable to consider the ternary complex as a six-state system in which each combination of the microstates of the two binary complexes exists. That is, the excited state of pMHC-II can exist with HLA-DM being bound via either the D1 subdomain alone, the D2 subdomain alone, or via both subdomains simultaneously. And, in principle, the same should hold for the ground state of pMHC-II. However, in order to explain the impact HLA-DM has on peptide dissociation, respectively the impact the peptide has on HLA-DM affinity, we have to establish an interaction between peptide and HLA-DM in the ternary complex. Given that HLA-DM and peptid do not directly interact with each other and the respective binding sites on the MHC-II molecule are non-overlapping (Fig. 1), this interaction is assumed to be allosteric and to result from opposing structural features of the two MHC-II conformations 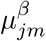 and 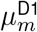. Specifically, we assume that 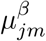 prohibits proper contact of HLA-DM to the D1 subdomain, and, conversely, MHC-II conformation 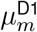 prevents the peptide from making proper contact with the β-sheet of the peptide binding groove. We refer to this as peptide-HLA-DM competition and take this into account by excluding the two corresponding microstates from the model. (This is again the above mentioned first defining property of optimal HLA-DM catalysis, and we will later relax this condition and discuss its implications in the section “Imperfect peptide-HLA-DM competition”.) Consequently, the ternary complex comprises four microstates (Fig. S3, ternary): microstate 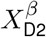 where the peptide-loaded binding groove is in its ground state while HLA-DM is bound only via D2, and the three microstates with the peptide-loaded binding groove in the excited state while HLA-DM is either bound only via the D2 subdomain,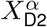, only via the D1 subdomain,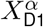, or via both subdomains simultaneously, 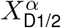. Given the spatial distance between the peptide binding groove and the D2 subdomain, we assume that both binding sites act independently. Consequently, the free energies of the four microstates are given by

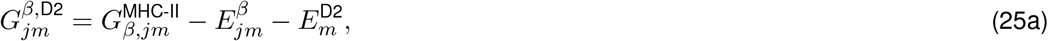

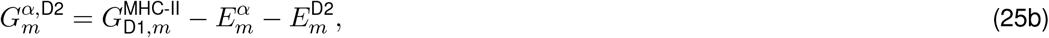

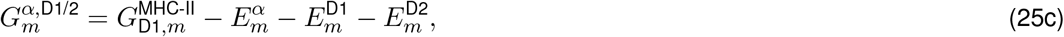

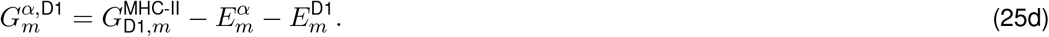

The equilibrium constants between the microstates depend on the free-energy differences of these states. For the transitions 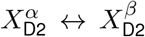 and 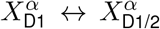 these are the same as their corresponding counterparts in the pMHC-II complex, respectively the MHC-II-HLA-DM complex (Fig. S3). The transition 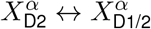 however, has a different equilibrium constant than the transition 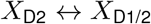, namely

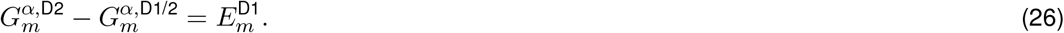

No reaction barrier hinders this transition anymore. The reason is that the excited state already comes with MHC-II conformation 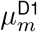, i.e., the excited state allows for efficient binding of HLA-DM to the D1 subdomain. This is one of the two effects of the above mentioned peptide-HLA-DM cooperativity; the second one is, as we will see below, that HLA-DM accelerates peptide association.

Having determined the free energies of the microstates of the ternary complex, we turn our attention to the population frequencies. On the population level, the concentration of the ternary complex relates to the concentrations of its microstates by

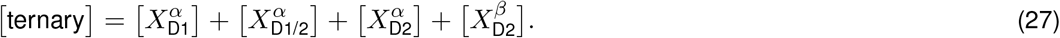

As before, we assume fast relaxation times on the molecular level. Thus, the population frequencies of the microstates of the ternary complex are

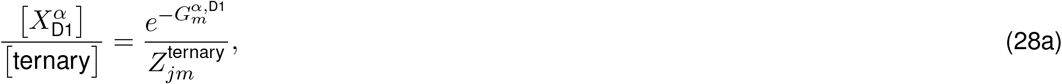

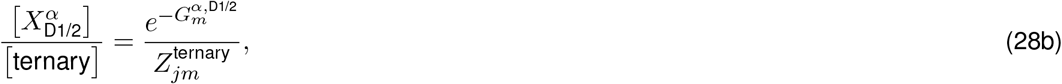

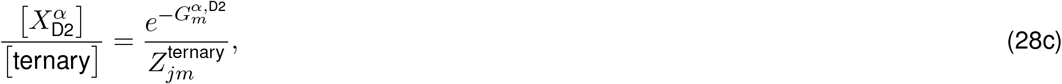

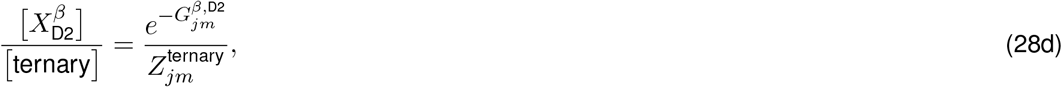

where we introduced the partition function of the ternary complex,

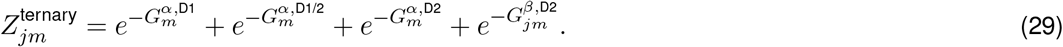

As before, relations (28) apply even when reactions are not (yet) in thermodynamic equilibrium on the population level.

Next, we explore the microscopic transitions that lead to the formation/decay of the ternary complex. We begin with the three microscopic transitions between the MHC-II-HLA-DM complex and the ternary complex. The first transition we consider is between the microstates *X*_D2_ and 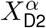. Since the peptide binding groove and the D2 subdomain are independent of each other, the microscopic kinetic rate constants are identical to those of the microscopic transition between microstate 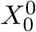 and *X*^*α*^ in the formation/decay of the pMHC-II complex. This can also be seen by comparing the free-energy difference of the microstates 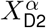 and *X*_D2_, Eqs. (25b) and (3a), to the free-energy difference of the microstates *X*^*α*^ and 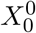, Eqs. (14b) and (1),

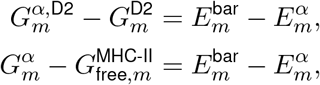

which are the same (and where we have used the definition of the reaction barrier, Eq. (6)). Next, we consider the transition between microstates *X*_D1/2_ and 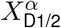. The free-energy difference between these microstates is

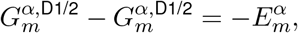

where Eqs. (25c) and (3b) were used. Thus, no reaction barrier is present anymore, because HLA-DM has removed it for the peptide. Consequently, this transition is no longer hindered by a suboptimally shaped MHC-II molecule. We assume that the entire effect of the removal of the reaction barrier manifests in the peptide association rate, which therefore is increased by a factor 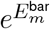 compared to the basal rate^1^.

As already pointed out above, this is the second effect of peptide-HLA-DM cooperativity—the peptide removes the reaction barrier for HLA-DM and HLA-DM removes the reaction barrier for the peptide. Lastly, we consider the transition between microstates *X*_D1_ and 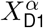. The free-energy difference between these microstates is

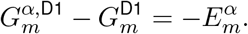

This is the same as for above transition, because the only difference between these two cases is whether HLA-DM is also bound to the D2 subdomain, which has no impact on the peptide binding groove. Thus, the kinetic parameters are the same as for the transition between microstates *X*_D1/2_ and 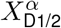.

Next, we consider the microscopic transitions between pMHC-II and the ternary complex. Two of these transitions account for binding/decay via the D2 subdomain. Due to the assumption that peptide binding groove and D2 subdomain are independent, the microscopic affinites for the transitions 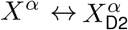 and 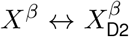 are the same and identical to the microscopic affinity of the transition 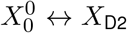, which is defined in Eq. (11b),

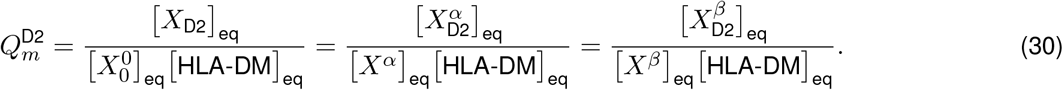

The third microscopic transition between pMHC-II and the ternary complex is via the D1 subdomain, 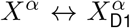. The microscopic affinity of this transition is not identical to the corresponding microscopic transition between microstates 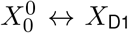, which is given by 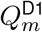, Eq. (11a). Instead, due to detailed balance, it is larger by a factor 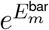.

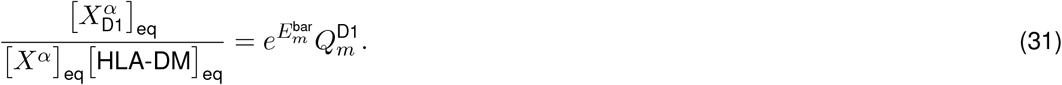

This is again a result of peptide-HLA-DM cooperativity, where the peptide has removed the reaction barrier HLA-DM is facing when binding to the D1 subdomain in the MHC-II-HLA-DM complex.

Since we are now familiar with the microscopic transitions between the two binary complexes and the ternary complex, we can proceed with deriving the macroscopic parameters. We begin with the peptide-modulated HLA-DM affinity. On the population level, this is defined by

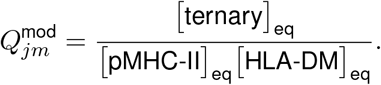

Using Eqs. (15)-(18) and (25)-(29), we arrive at

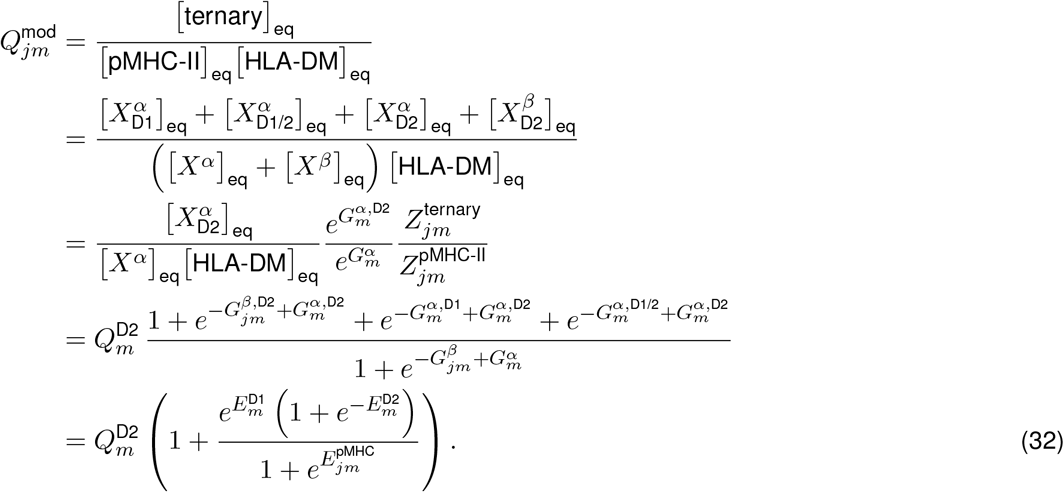

Next, we derive the catalyzed binding kinetics of the peptide. We consider the rate of change of the MHC-II-HLA-DM complex. Since we are here not interested in actually solving the system but merely want to derive expressions for the macroscopic rate constants by comparison of terms, we ignore reaction fluxes between free MHC-II and the MHC-II-HLA-DM complex. It holds

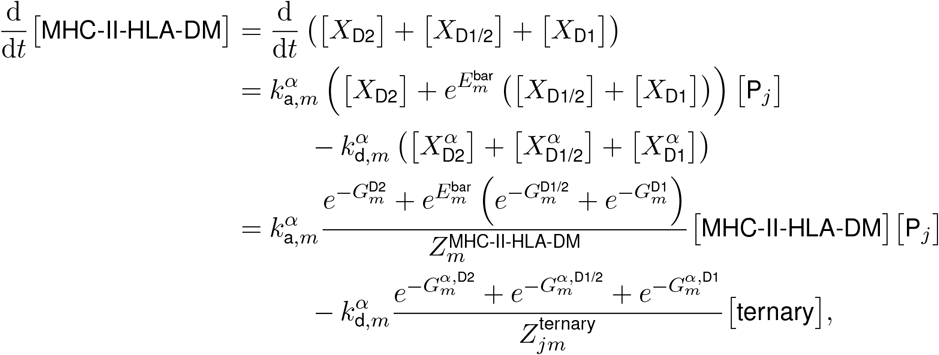

and by comparing this to

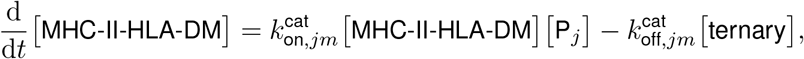

we find

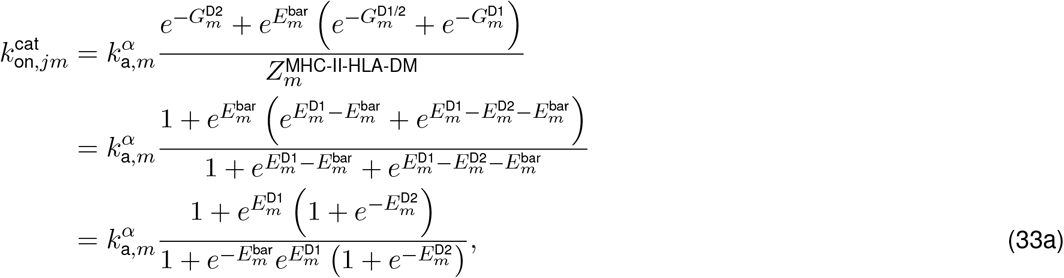

and

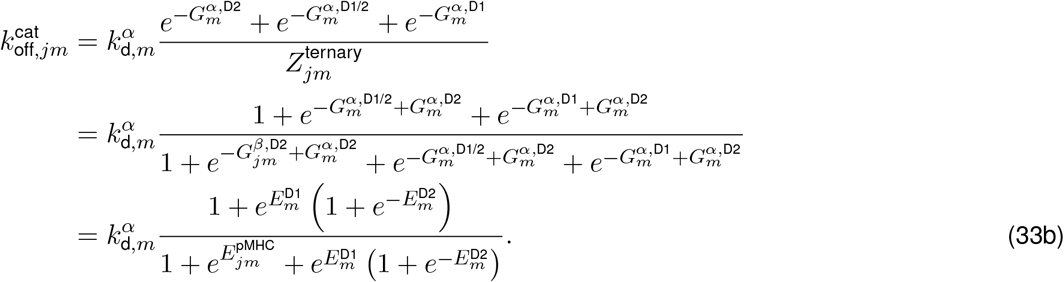

### Observational constraints on the microscopic model

In the previous section, we derived equations for the macroscopic parameters of the stoichiometry-based model in terms of the microscopic parameters of the structure-based model, which are given in Eqs. (13), (22), (32), and (33). These equations depend in total on six peptide-unspecific parameters: 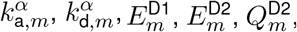, and 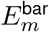. Upon closer inspection of these equations, we note that neither 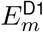 nor 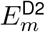 appear separated from each other in any of the equations, but only as the combination 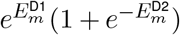. This means that neither of these parameters can be estimated individually from the data. To account for this, we introduce an effective parameter, the energetic potential of HLA-DM binding,

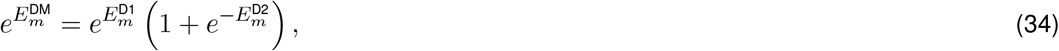

which reduces the number of peptide-unspecific parameters from six to five. The implication of this reduction is that the two microstates with HLA-DM bound by the D1 subdomain, i.e., *X*_D1/2_ and *X*_D1_ in the MHC-II-HLA-DM complex and 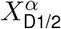 and 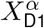 in the ternary complex, become effectively indistinguishable from each other—they behave as a single, effective microstate. Thus, the MHC-II-HLA-DM complex is only “seen” as a two-state system under the experiment, which also applies to HLA-DM binding in the ternary complex. Moreover, we can no longer resolve the detailed-balance relation (12), since this requires knowledge of 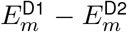, which we cannot obtain from the data. The consequence of this is that we are unable to make any quantitative statement about the microscopic affinity 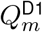, defined in Eq. (11a). Taken together, even though we describe HLA-DM binding to the D1 and D2 subdomain by three microstates and allow for formation and decay by either subdomain, the data are only informative about two microstates and the thermodynamics of formation and decay via the D2 subdomain. The model shown in Fig. 4A has taken these observational constraints into account, where we denote the merged, effective microstates with *X*_D1_ (MHC-II-HLA-DM complex) and 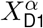 (ternary complex). Furthermore, we use 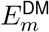 in the equilibrium constants of the transitions *X*^*α*^ *↔ X*_D1_ and 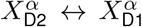 in the effective model, and we omit HLA-DM binding to the D1 subdomain from solution.

We now have to deal with another observational constraint, which is concerned with the fact that the variable factor in the experimental data is the nature of the peptide. The only peptide-specific parameter in the equations is 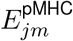, which, mathematically, acts as the independent variable. In contrast, the data allow only for comparisons with the basal peptide dissociation rate—no direct information about the pMHC-II binding energy are available. To account for this, we have to transform our equations such that 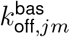 can act as independent variable. In order to do this, we note that in any of our equations in which the pMHC-II binding energy appears, it does so in the form 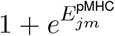. From Eq. (22b) we see that this relates to 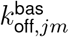 by

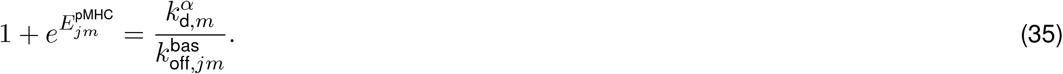

Using this together with constraint (34), we obtain the expressions for the macroscopic parameters that can be compared to population-level data,

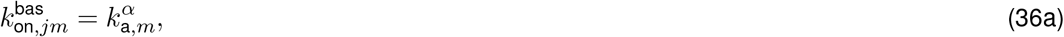

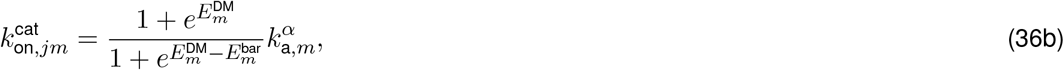

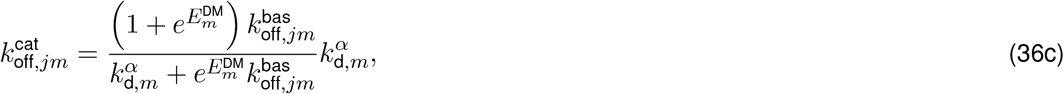

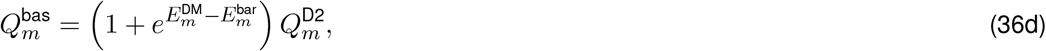

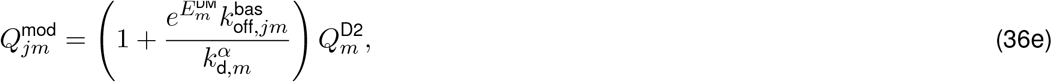

which contain five peptide-unspecific parameters and have the basal peptide dissociation rate as independent variable. The data shown in Fig. 2 only include direct information about 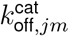 and 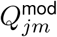 (and further macroscopic parameters that can be derived from them, Fig. 2C). These two functions depend on only three of the five peptide-unspecific parameters: 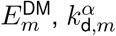, and 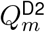. In order to determine the remaining two, 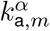 and 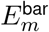, association data for a comprehensive peptide set are required.

The three peptide-unspecific parameters we can get our hands on using available data appear as asymptotic values of the equations. It holds

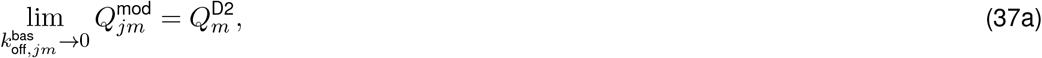

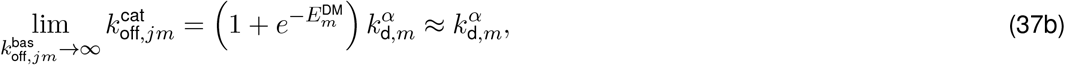

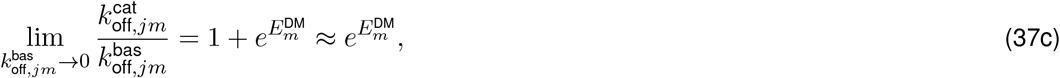

where we have used 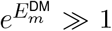 in the approximations, which applies if the energetic HLA-DM potential lies well above the thermal energy. Thus, these values can be read off the data. Given the substantial amount of variation in the data that is not explained by this single-variable version of our model, it is a priori not clear how to estimate these values and whether those estimates are meaningful. In the face of the model extensions discussed in the next sections, however, it turns out that this model version should be regarded as a limiting case of the more general model. Specifically, this limiting case describes optimal HLA-DM catalysis, which applies to peptides against which HLA-DM can most potently act, see the next section. The values stated in Fig. 3H were chosen to meet this criterion.

### Variations beyond optimal HLA-DM catalysis

In our structure-based model, HLA-DM exerts its impact on peptide-MHC-II interactions by two allosteric effects. Firstly, when entering and leaving the peptide binding groove, the peptide induces an MHC-II conformation,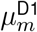, that enables optimal HLA-DM binding with the D1 subdomain, and likewise, when HLA-DM induces this MHC-II conformation when in the MHC-II-HLA-DM complex, the peptide can enter the peptide binding groove much faster, because the reaction barrier the peptide faces in the absence of HLA-DM has been removed. We refer to this as peptide-HLA-DM cooperativity. Secondly, the MHC-II conformation of the pMHC-II ground state,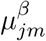, perfectly excludes HLA-DM binding to the D1 subdomain, resulting in HLA-DM accelerating peptide dissociation in a pMHC-II-binding-energy-dependent manner. We refer to this as peptide-HLA-DM competition.

The microscopic model discussed in the previous section describes optimal HLA-DM catalysis, i.e., it describes both optimal peptide-HLA-DM cooperativity and optimal peptide-HLA-DM competition. Optimal peptide-HLA-DM cooperativity refers to the MHC-II conformation of the excited state of the pMHC-II complex, respectively of microstate 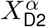 of the ternary complex. The best possible MHC-II conformation for HLA-DM binding that the peptide can induce is the one that HLA-DM induces itself in the MHC-II-HLA-DM complex, i.e., MHC-II conformation 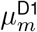 (Fig. S3, light blue MHC-II conformation). However, if the peptide side chains play a role in the transition 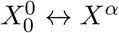, respectively in the transition 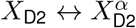, the MHC-II conformation the peptide induces in the excited state may deviate from 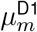 in a peptide-specific manner. Moreover, the binding energy in the excited state will then in general depend on the nature of the peptide as well, which may further counteract HLA-DM’s ability to force the ternary complex into microstate 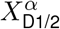 (or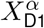), i.e., to destabilize the peptide. This is the basis for imperfect peptide-HLA-DM cooperativity and is discussed in the next section.

Optimal peptide-HLA-DM competition refers to the complete inability of HLA-DM to bind to the D1 subdomain as long as the MHC-II-bound peptide resides in the ground state. The competition between peptide and HLA-DM is assumed to be mediated by key amino acids of the MHC-II molecule to be constantly inward-flipped (or otherwise out of reach for the complementary anchor points on the HLA-DM molecule) in microstates *X*^*β*^, respectively 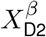. However, molecular dynamics simulations suggest that those elements can spontaneously flip outwards even in the ground state of the pMHC-II complex [7]. The frequency with which these spontaneous flipping events occur may be dependent on the nature of the peptide—for some this may occur more frequently than for others. This implies imperfect peptide-HLA-DM competition.

We first discuss imperfect peptide-HLA-DM cooperativity, because, as the analysis reveals, it is the cause behind highly HLA-DM resistant peptides, and it is what the peptide-specific parameter *x*_*jm*_ of the model shown in Fig. 4A effectively describes. We then proceed to imperfect peptide-HLA-DM competition, where we will see that it has only minor impact on the macroscopic stoichiometry-based parameters.

### Imperfect peptide-HLA-DM cooperativity

As pointed out above, imperfect HLA-DM cooperativity relates to the MHC-II conformation in the excited state of pMHC-II (Fig. S4). We no longer require this MHC-II conformation to be 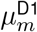, and we denote the peptide-specific MHC-II conformation the peptide induces in the excited state with 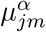 ((Fig. S4, purple MHC-II conformation), and the corresponding free energy of the MHC-II molecule is 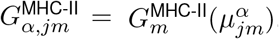. Furthermore, the binding energy in the excited state between the peptide and the binding groove of the MHC-II molecule is then in general peptide-specific as well; we denote it with 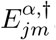. Thus, the free energy of the excited state now reads

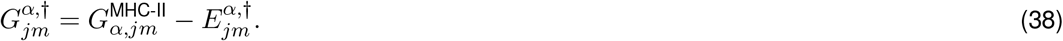

It can be expected that this peptide-specific excited state only plays a role for peptides for which 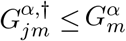, since the pathway through MHC-II conformation 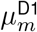 would otherwise offer an alternative of lower free energy. The free-energy difference between the maximally cooperative excited state, i.e., the one of the previous sections that has the MHC-II molecule in conformation 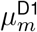, and the imperfectly cooperative excited state, i.e., the one we are discussing in this section, is denoted

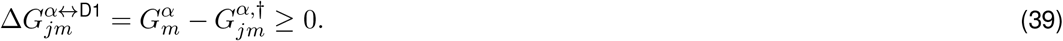

Since the free energy of the excited state is now peptide-specific, the two microscopic kinetic rate constant of the transition 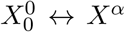 must be considered peptide-specific as well. We take this into account by introducing two new parameters, *x*_a,*jm*_ and *x*_d,*jm*_, that account for the peptide-specific variations in the microscopic association and dissociation rates (Fig. S4, highlighted in red). With this, we can derive the macroscopic parameters, which otherwise exactly parallels the calculation in the section on the pMHC-II complex, and we find

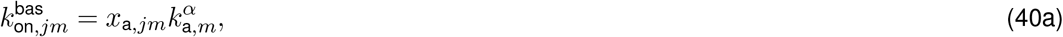

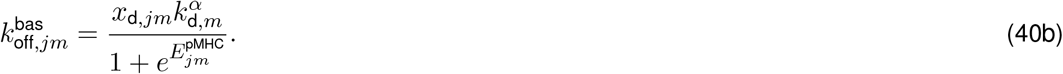

We note that basal peptide affinity now reads

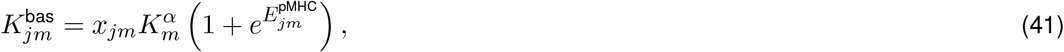

where we introduced the ratio of the two variation parameters,

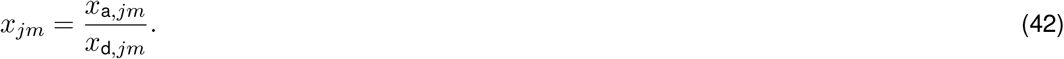

While *x*_a,*jm*_ and *x*_d,*jm*_ are kinetic parameters, *x*_*jm*_ determines the thermodynamics of imperfect peptide HLA-DM cooperativity.

The modifications to the excited state also leave their mark in the ternary complex. Specifically, the free energy of microstate 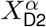, Eq. (25b), must be adjusted to

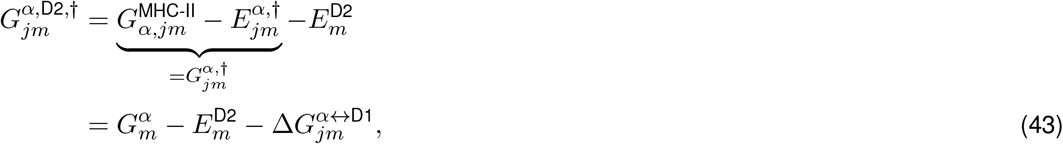

which, in the second step, we express in terms of the free energy of the excited state for perfect peptide-HLA-DM cooperativity, 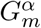, and the free-energy difference 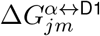 defined in Eq. (39). The remaining free energies of the ternary complex, Eqs. (25), stay unchanged. Consequently, the kinetic parameters of the microscopic transition 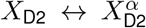 are subject to the same modifications as their counterpart 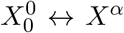, i.e., they are equipped with the variation parameters *x*_a,*jm*_ and *x*_d,*jm*_ (Fig. S4, highlighted in red). Furthermore, detailed balance implies that the free-energy difference 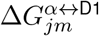 is related to the parameter *x*_*jm*_, defined in Eq. (42), by

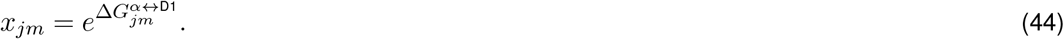

The partition function of the ternary complex becomes

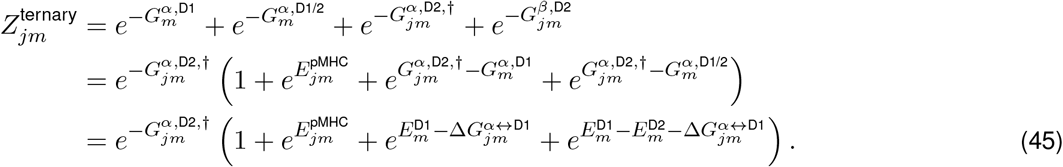

With this we can use the same steps as in the section on the ternary complex to arrive at expressions for peptide-modulated HLA-DM affinity and catalyzed peptide association and dissociation rates,

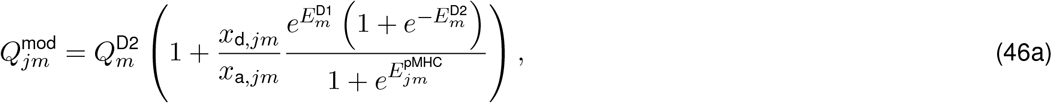

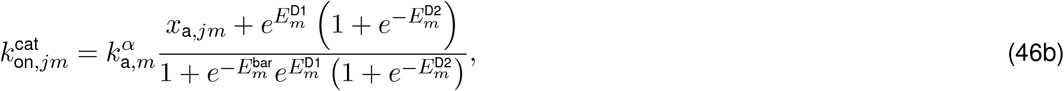

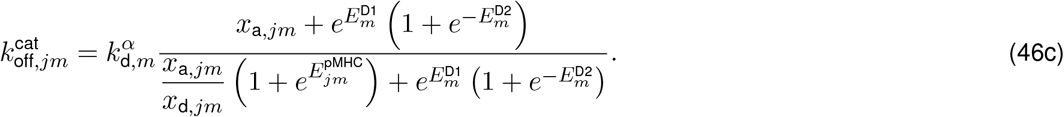

Eqs. (40) and (46) depend on the same six peptide-unspecific parameters as the equations for optimal HLA-DM catalysis, but they depend on three peptide-specific parameters, the pMHC-II binding energy 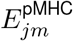, the variation parameter for basal peptide association *x*_a,*jm*_, and the variation parameter for microscopic dissociation from the excited state, *x*_d,*jm*_. We will now bring these equations into the form that meets the observational constraints of the experimental data.

The equations have precisely the same properties regarding the parameters 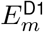 and 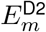 as discussed in Section “Observational constraints on the microscopic model”, including all the mentioned limitations. In particular, we have to restrict ourselves to using the energetic potential of HLA-DM binding, Eq. (34). What is left to do is to make basal peptide dissociation an independent variable of the equations. From Eq. (40b), we see that

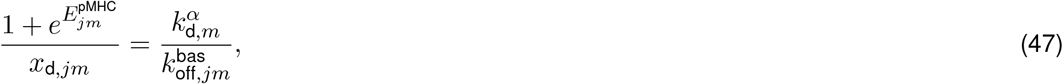

where the right-hand-side contains only the single peptide-specific parameter 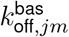, while the left-handside contains two peptide-specific parameters, namely 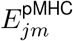 and *x*_d,*jm*_. Importantly, this precise combination of these two parameters also appears in the expressions for peptide-modulated HLA-DM affinity, Eq. (46a), and catalyzed peptide dissociation rate, Eq. (46c). Inserting Eq. (47) into these two expressions leads to

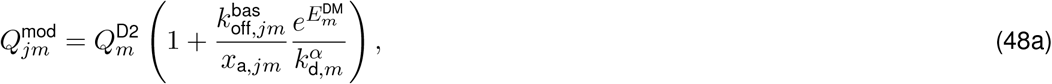

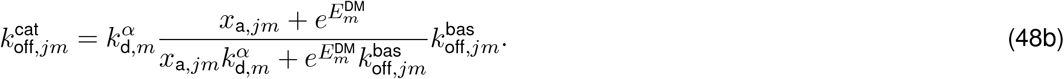

Surprisingly, the final equations depend on only two peptide-specific parameters: the basal peptide dissociation rate 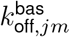 and the variation parameter of basal peptide association rate *x*_a,*jm*_. The interpretation of this is that the observations on the population level may be able to determine the value of the basal peptide dissociation rate but they do not reveal the microscopic details of how this value comes about— two peptides that differ in their values for *x*_d,*jm*_ and 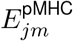 but have the same 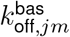 are indistinguishable from each other on the population level, provided they also have the same 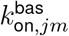.

Given that Eqs. (39) and (42) imply that *x*_*jm*_ *≥* 1, we expect that in general *x*_a,*jm*_ *≥* 1. Then, the case *x*_a,*jm*_ = 1 does describe peptides against which HLA-DM is the most potent, justifying the notion of optimal HLA-DM catalysis and the corresponding criterion of estimating the peptide-unspecific parameters from the data.

### Imperfect peptide-HLA-DM competition

When discussing the ternary complex for optimal HLA-DM catalysis, we neglected two microstates based on the assumption that simultaneous contacts of peptide with the β-sheet and of HLA-DM with the D1 subdomain are completely impossible, establishing optimal peptide-HLA-DM competition. Now, we are going to relax this condition and discuss the imperfect case by reinstating these two microstates, which are denoted 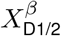 and 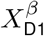 (Fig. S5). In order to maintain peptide-HLA-DM competition, we introduce a competition term 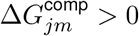 into the corresponding free energies,

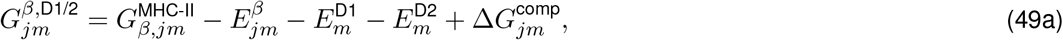

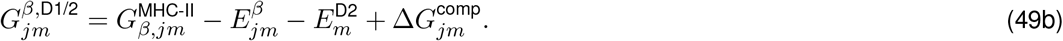

The remaining free energies stay the same as in Eqs. (25). The partition function is then

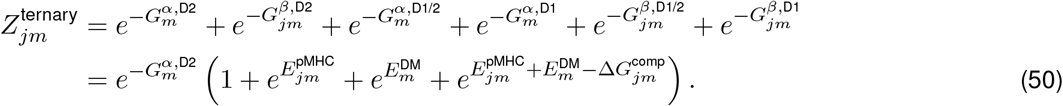

Defining

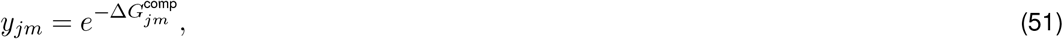

we follow the same steps as in the section on the ternary complex to derive the expressions of peptide-modulated HLA-DM and catalyzed peptide dissociation rate,

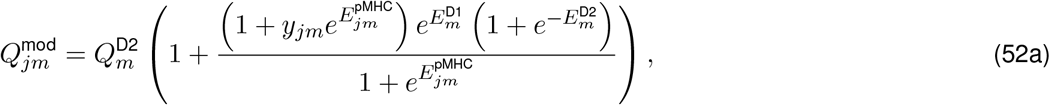

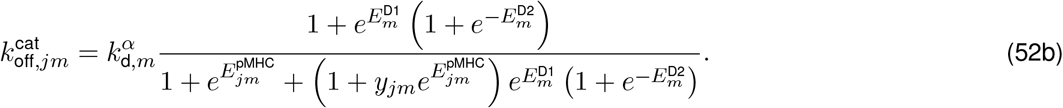

All other equations are the same as for optimal HLA-DM catalysis. Thus, the equations for imperfect peptide-HLA-DM competition also depend on the same six peptide-unspecific parameters as optimal HLA-DM catalysis. As before, the equations have the same dependence on parameters 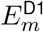 and 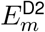, bearing the same consequences, and the energetic potential of HLA-DM binding 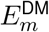, defined in Eq. (34), has to be used in the final equations. Next to the pMHC-II binding energy 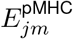, the equations depend on the additional peptide-specific parameter *y*_*jm*_. In order to make the basal peptide dissociation rate an independent variable, we note from Eq. (22b) that

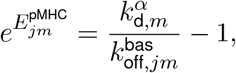

and we find

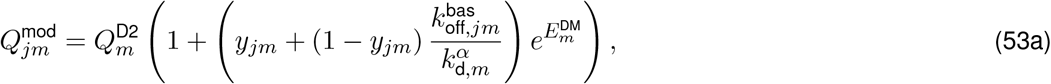

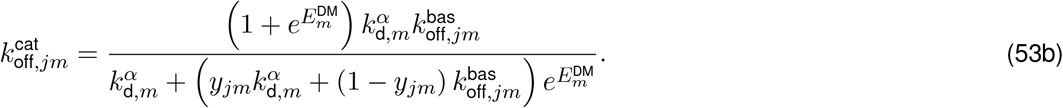

The equations for optimal HLA-DM catalysis are recovered for *y*_*jm*_ = 0. The case *y*_*jm*_ = 1 describes the complete absence of peptide-HLA-DM competition, where peptide and HLA-DM can optimally bind to MHC-II without interfering with each other. In the following, we assume that there is always competition between peptide and HLA-DM, it just does not have to be perfect: 0 *≤ y*_*jm*_ *«* 1.

Imperfect peptide-HLA-DM competition manifests differently for weakly and strongly binding peptide. For weakly binding peptides, imperfect peptide-HLA-DM competition becomes negligible, because

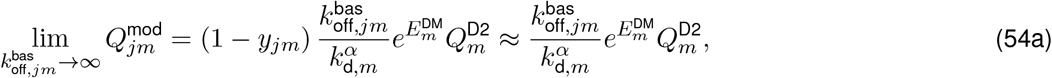

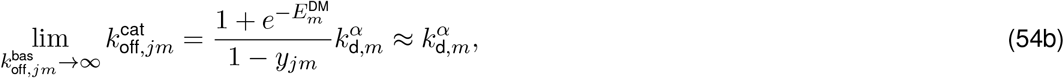

which, since 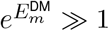 and *y*_*jm*_ *«* 1, do not depend on *y*_*jm*_. However, for strongly binding peptides we find

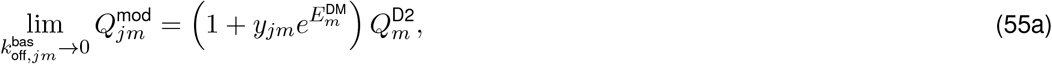

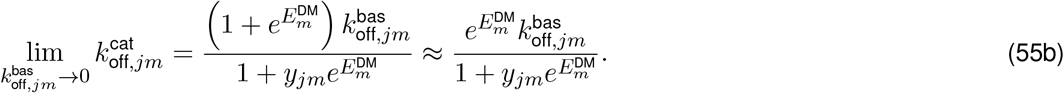

Thus, peptide-HLA-DM competition is practically perfect if 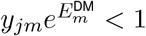, which, using Eq. (51), translates to 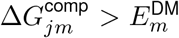. For peptides for which 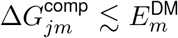, observable effects manifest as variation of peptide-modulated HLA-DM affinity and catalyzed peptide dissociation rate for strongly binding peptides. This is similar for HLA-DM impact on dissociation. Since imperfect peptide-HLA-DM competition on its own has no impact on basal peptide association rate, HLA-DM susceptibility and relative HLA-DM susceptibility are not affected by the variable *y*_*jm*_. An important consequence of this is that *y*_*jm*_ plays only a role at rather high HLA-DM concentrations because HLA-DM-dependent peptide dissociation is governed by susceptibility at low HLA-DM concentrations.

**Figure S1:**
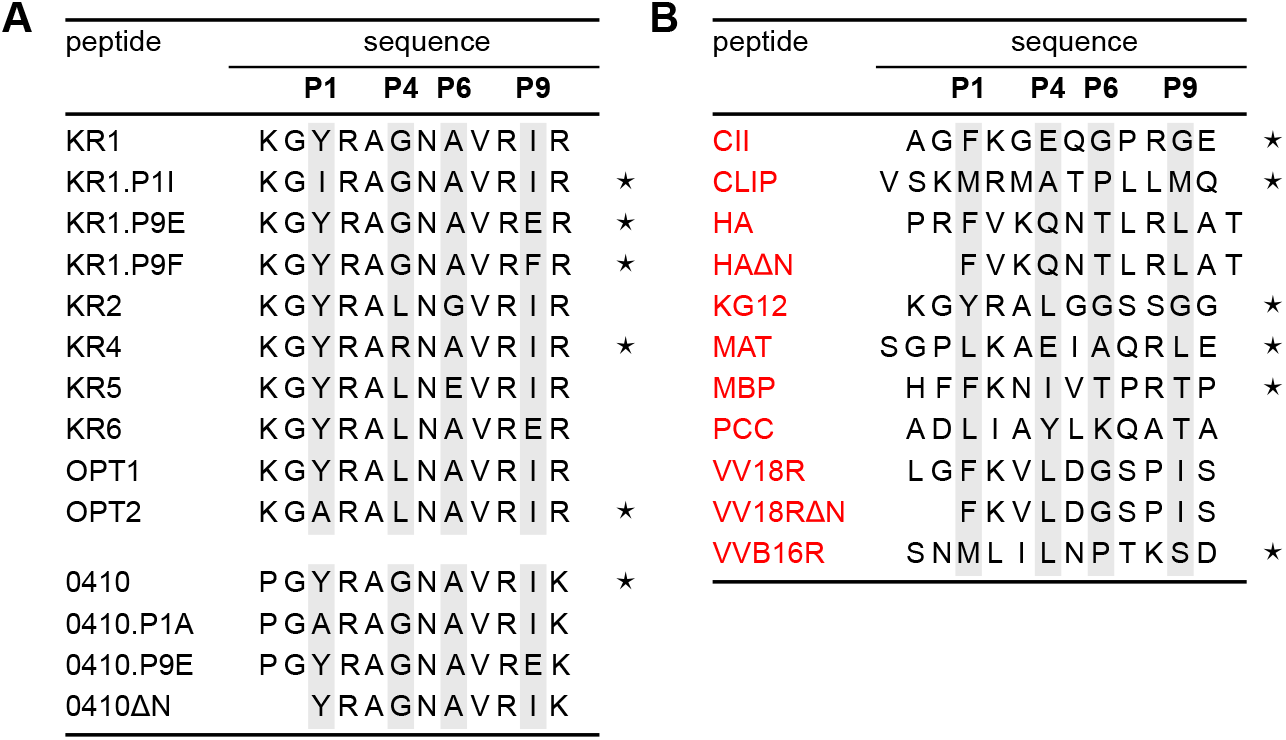
Peptide set contains closely related sequences and unrelated sequences. (A) The two groups of related peptide sequences. In each group variations exist only in the main anchor points (gray highlighting). (B) Sequences of unrelated peptides. Peptides with common catalyzed peptide dissociation rate, gray area in Fig. 2E, are marked by a star.

**Figure S2:**
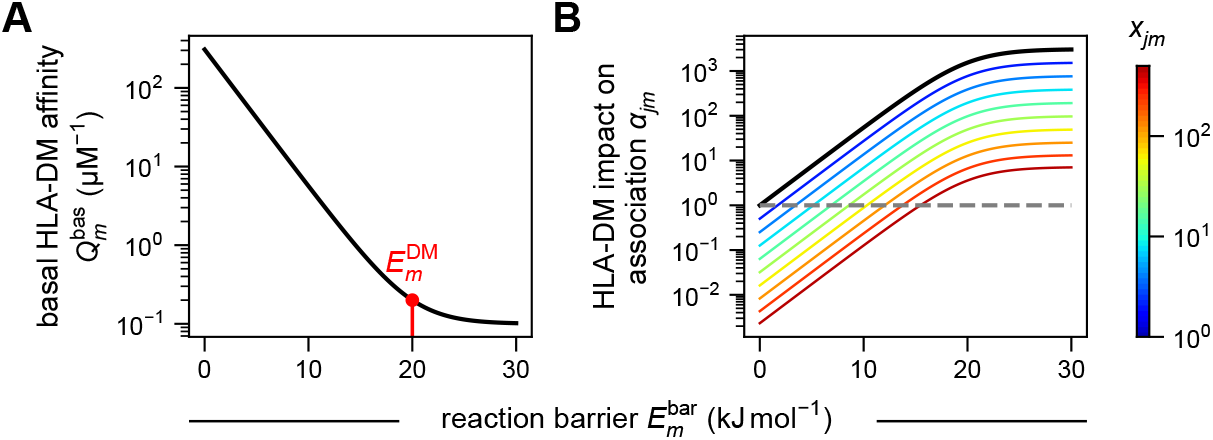
Low basal HLA-DM affinity is a prerequisite for HLA-DM accelerating peptide association. (A) Basal HLA-DM affinity as a function of the reaction barrier. (B) HLA-DM impact on association as a function of the reaction barrier for peptides of various basal peptide association rates, parameterized by *x*_*jm*_ (color bar). The black curve corresponds to *x*_*jm*_ = 1, and the gray dashed curve indicates *α*_*jm*_ = 1.

**Figure S3:**
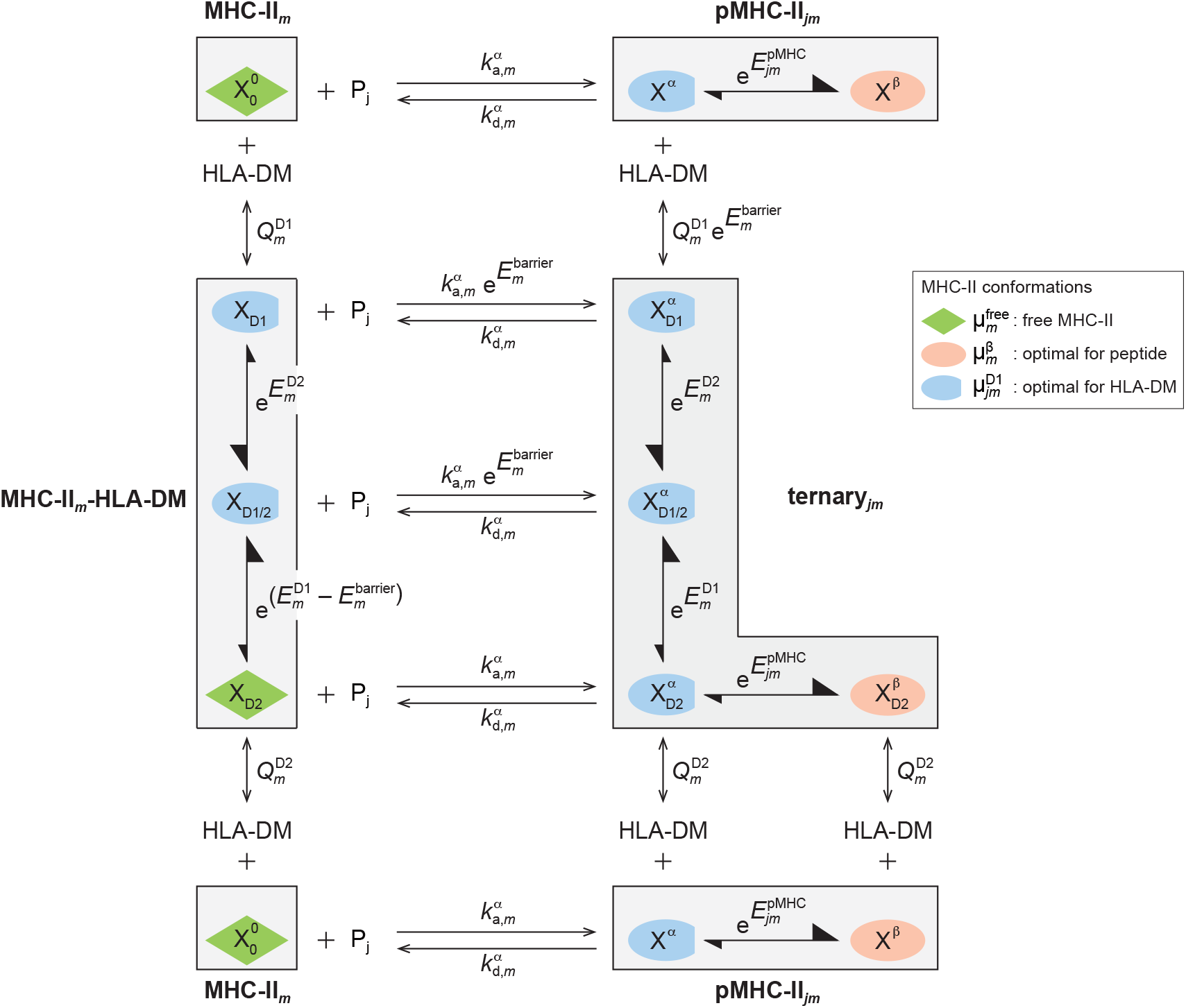
Structure-based model of optimal HLA-DM catalysis.

**Figure S4:**
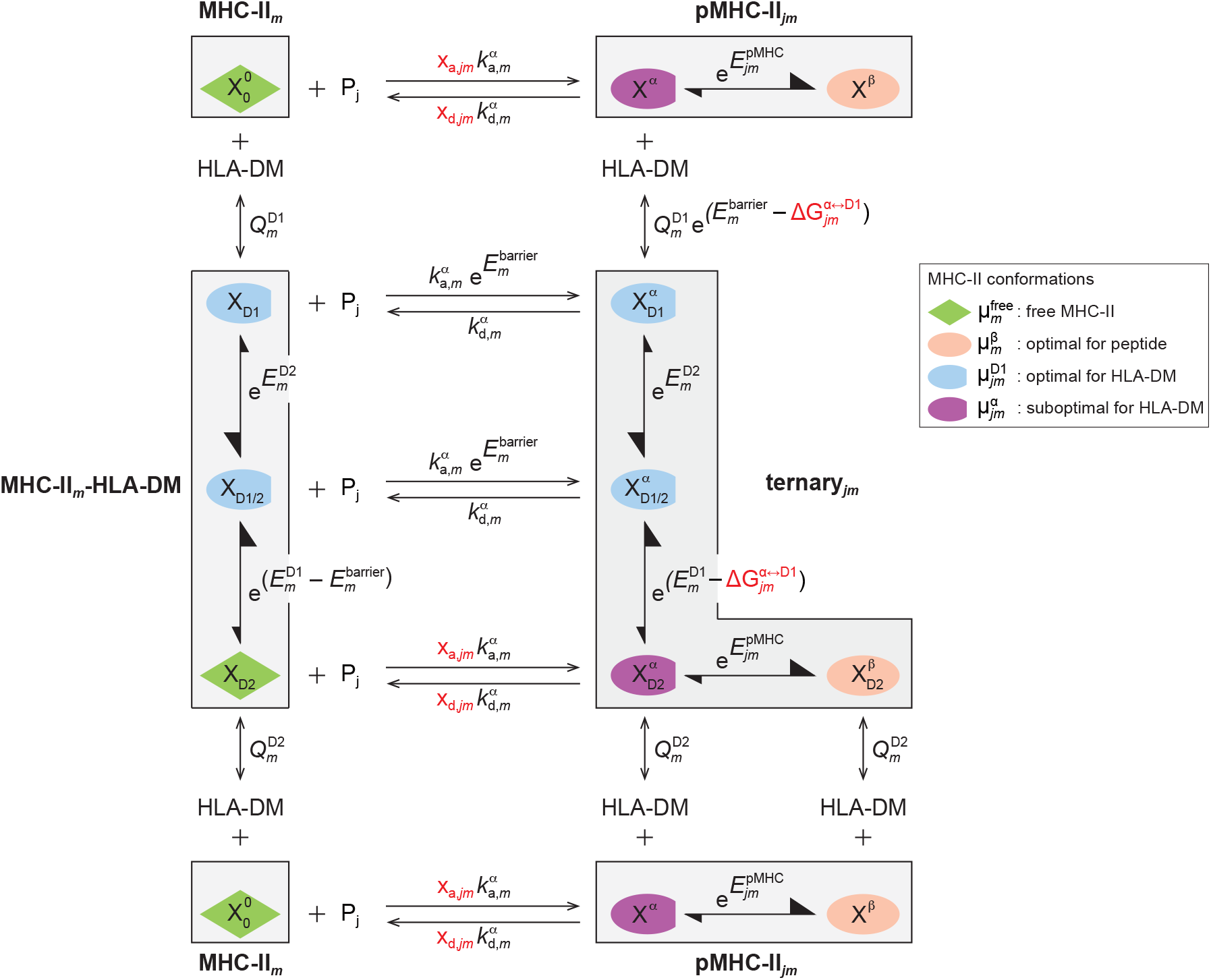
Structure-based model of imperfect peptide-HLA-DM cooperativity.

**Figure S5:**
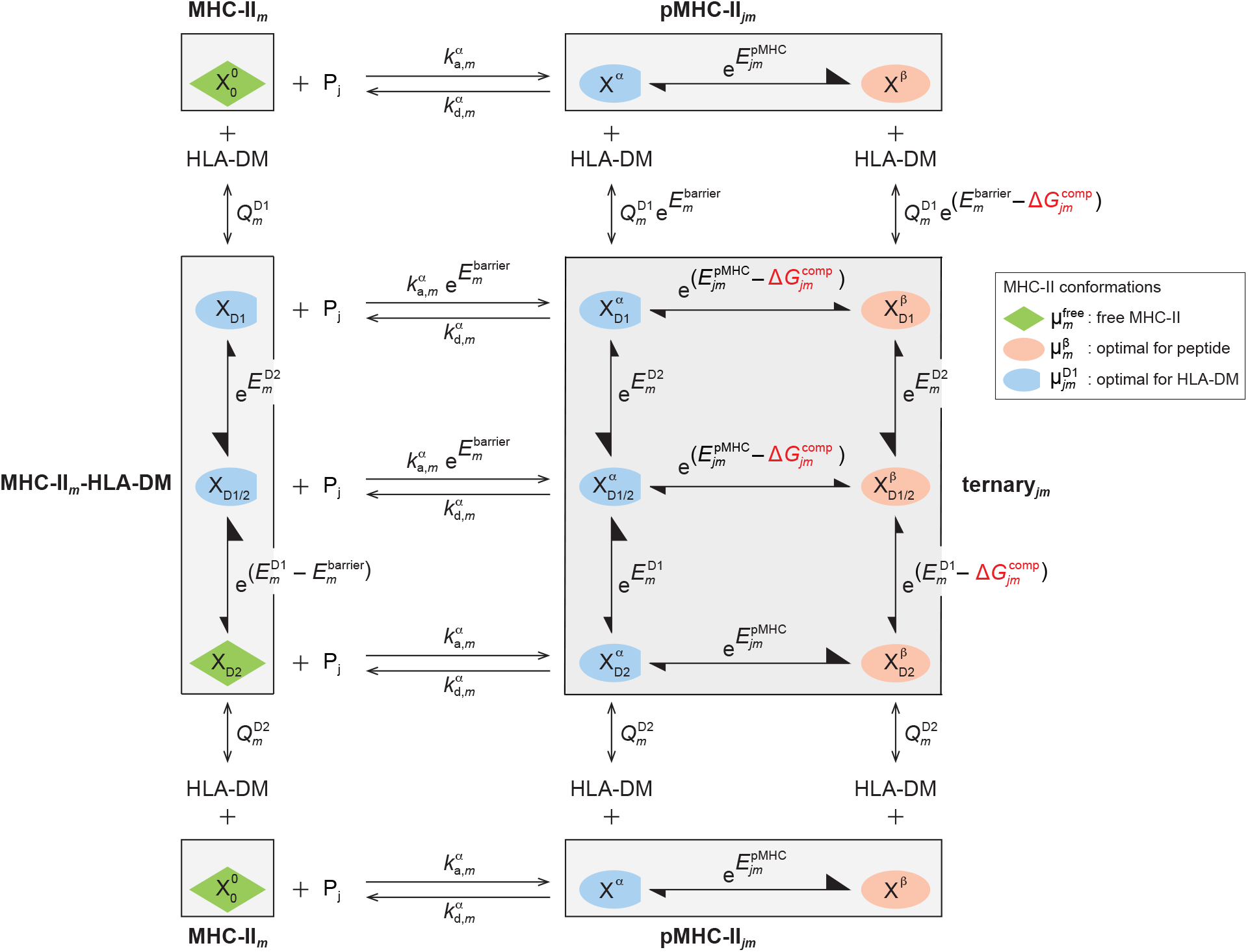
Structure-based model of imperfect peptide-HLA-DM competition.

## Footnotes

1 Technically, this does not have to be the case, and only part of the removal may manifest in peptide association, while the remaining part manifests in the peptide dissociation rate from microstate 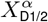. This can in principle be determined based on fluorescence anisotropy measurements, but this requires association data from a comprehensive set of peptides measured in the context of the same allotype. To our knowledge, such a data set does not exist. Since there is nevertheless ample evidence that HLA-DM does accelerate peptide association, we decided to shift the entire effect of reaction-barrier removal into the association rate, and leave it to experimentalists to provide the necessary data that allow to refine this detail in our model.

